# Glycolysis-enhancing α1-adrenergic antagonists are neuroprotective in Alzheimer’s disease

**DOI:** 10.1101/2025.04.03.647018

**Authors:** Qiang Zhang, Jordan Schultz, Jacob Simmering, Braedon Q Kirkpatrick, Matthew A Weber, Sydney Skuodas, Tara Hicks, Grace Pierce, Margaret Laughlin, Aimee X Bertolli, Travis Larson, Ramasamy Thangavel, Mayu Oya, David Meyerholz, Georgina Aldridge, Jan Fassler, Nandakumar S. Narayanan

## Abstract

Terazosin (TZ) is an α_1_-adrenergic receptor antagonist that enhances glycolysis by activating the enzyme phosphoglycerate kinase 1 (PGK1). Epidemiological data suggest that TZ may be neuroprotective in Parkinson’s disease and in dementia with Lewy bodies and that glycolysis-enhancing drugs might be protective in other neurodegenerative diseases involving protein aggregation, such as Alzheimer’s disease (AD). We investigated TZ in AD and report four main results. First, we found that TZ increased ATP levels in a *Saccharomyces cerevisiae* mutant with impaired energy homeostasis and reduced the aggregation of the AD-associated protein, amyloid beta (Aβ) 42. Second, in an AD transgenic mouse model (5xFAD) we found that TZ attenuated amyloid pathology in the hippocampus and rescued cognitive impairments in spatial memory and interval timing behavioral assays. Third, using the Alzheimer’s Disease Neuroimaging Initiative (ADNI) database, we found that AD patients newly started on TZ or related glycolysis-enhancing drugs had a slower progression of both cognitive dysfunction and neuroimaging biomarkers, such as ^18^F-fluorodeoxyglucose positron emission tomography (FDG-PET), a measure of brain metabolism. Finally, in a large human administrative dataset, we found that patients taking TZ or related glycolysis-enhancing drugs had a lower hazard of being diagnosed with AD compared to those taking tamsulosin or 5-alpha reductase inhibitors. These data further implicate metabolism in neurodegenerative diseases and suggest that glycolysis-enhancing drugs may be neuroprotective in AD.

## INTRODUCTION

Alzheimer’s disease (AD) is the leading cause of dementia and affects over 6 million Americans (Rajan et al., 2021). The pathophysiology of AD involves complex mechanisms, including amyloid-beta (Aβ) aggregation, tau pathology, and neuronal loss, which cause synaptic dysfunction and cognitive impairment. Despite recent advances (Mintun et al., 2021; van Dyck et al., 2023), there is an ongoing need for better therapies for AD.

A factor that may contribute to AD is impaired energy metabolism (Mosconi, 2005a, 2005b; Profenno et al., 2010). Four lines of evidence support this link: 1) Aging is a primary risk factor for AD as it associated with impaired cerebral glucose metabolism, reduced mitochondrial biogenesis, and decreased adenosine 5′-triphosphate (ATP) levels (Herrup, 2010; Saxena, 2012). 2) Diseases of metabolism such as diabetes and obesity increase the risk of AD (Mosconi, 2005a; Profenno et al., 2010). 3) Interventions that improve metabolism, such as caloric restriction and exercise, can improve neuropathology and behavior in mouse models of AD and cognitive function in humans with AD (Intlekofer & Cotman, 2013; Wang et al., 2005; Witte et al., 2009). 4) Deficits in glucose metabolism can diagnose AD using ^18^F-fluorodeoxyglucose positron emission tomography (FDG-PET), as glycolysis is reduced in brain areas with amyloid deposition and tau accumulation (Minoshima et al., 1995; Vlassenko et al., 2010; Vlassenko & Raichle, 2015). These data lead to the hypothesis that enhancing glycolysis is neuroprotective in AD.

We tested this hypothesis by investigating the glycolysis-enhancing drug terazosin (TZ) in AD. TZ is widely used as an alpha-1 (α_1_) adrenergic receptor antagonist for hypertension and benign prostatic hyperplasia; however, it has another target: phosphoglycerate kinase 1 (PGK1), the first ATP-generating step that enhances glycolysis. TZ increases energy metabolism in animal models of Parkinson’s disease (PD) and in patients with PD (Cai et al., 2019a; Schultz et al., 2022). TZ is neuroprotective in animal models of PD (Cai et al., 2019b; Weber et al., 2023). It is also associated with slower disease progression in patients with PD and a decreased risk of being diagnosed with PD in humans (Cai et al., 2019b; Simmering et al., 2021, 2022), and mitigates neurodegeneration in the APPsw/PS1 transgenic mouse model of AD (Yu et al., 2022). In the present study, we investigated the use of TZ in yeast cellular models, in a mouse model of AD, in humans from the Alzheimer’s Disease Neuroimaging Initiative (ADNI) cohort, and in a large database of health insurance claims. These findings contribute to the growing body of evidence focused on metabolism in neurodegenerative disease and highlight the potential of glycolysis-enhancing drugs as a treatment for patients with AD.

## RESULTS

### TZ increases ATP in yeast and mitigates amyloid beta (Aβ) 42 aggregation

We investigated the effects of TZ in yeast, a eukaryotic organism with a glycolytic pathway that is conserved in humans. To investigate the effects of TZ on cellular energetics (Imamura et al., 2009; Nakano et al., 2011; Takaine, 2019a; Takaine et al., 2019) and Aβ42 aggregation, we quantified ATP at the single cell level using a fluorescent ATP biosensor (QUEEN, quantitative evaluator of cellular energy) (Takaine, 2019b; Yaginuma et al., 2014) and at the population level using a luciferase-based biochemical assay. ATP levels were lower in *snf1* mutant yeast strains compared to wild-type (WT) strains (Fig 1A–B; p = 0.0001 QUEEN; p = 0.03 luciferase) as expected (Hedbacker, 2008). Using a biochemical assay, we validated the reduction in ATP and determined that the *snf1* mutant cells had an average ATP concentration of 1.2 mM, which was 40% of the 3.7 mM ATP levels we observed in WT cells (Fig. 1B).

**Figure. 1.**
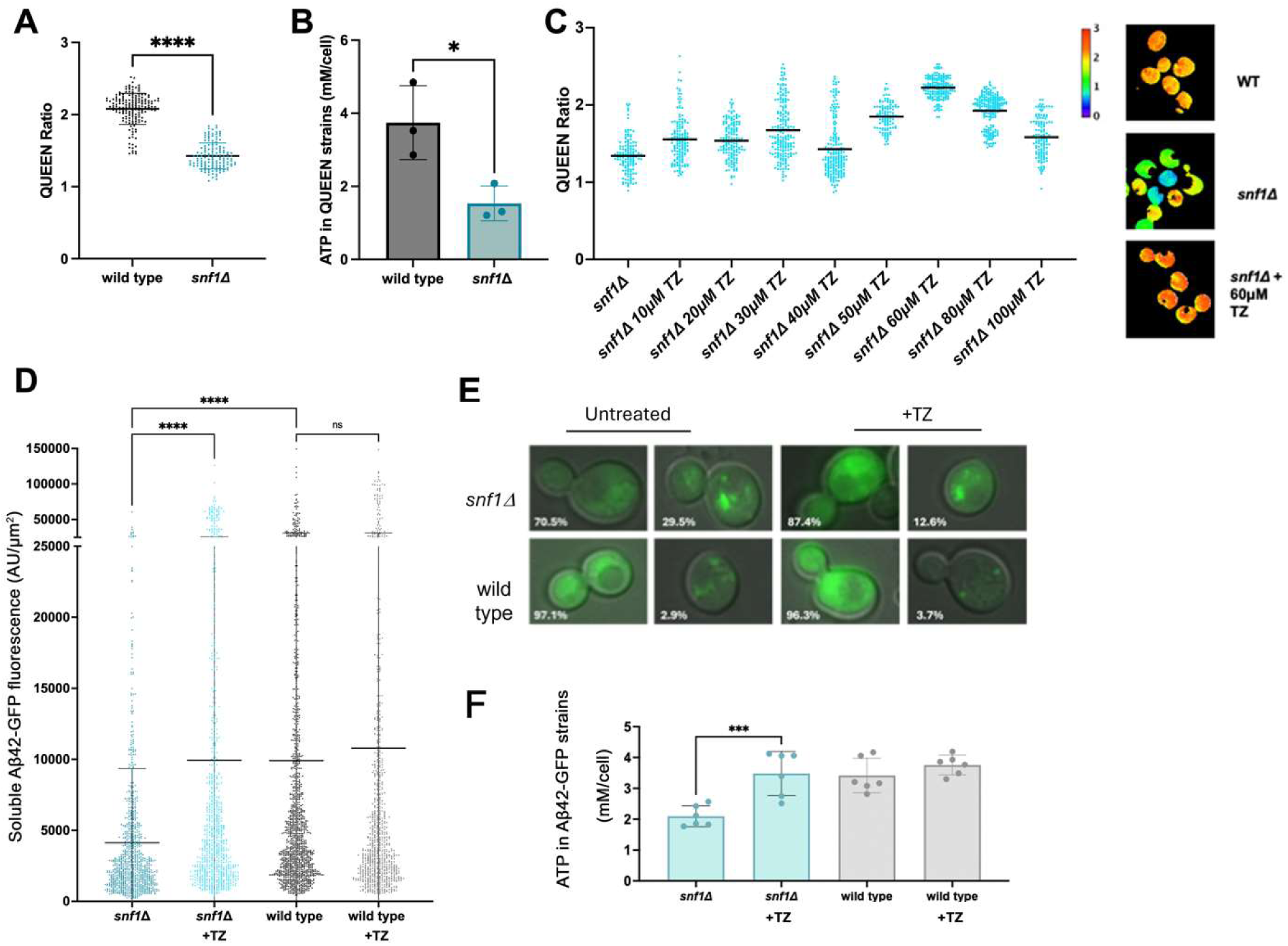
Terazosin (TZ) increases ATP levels and mitigates Aβ42-GFP aggregation in yeast. A) A reduction in ATP levels in the *snf1* mutant compared to wild-type yeast strains based on the QUEEN ATP biosensor. Each QUEEN data point represents the ratio within an individual cell; n=500 cells per dose. B) Luciferase-based biochemical measurement of cellular ATP levels conducted in triplicate (*p=0.03). C) ATP dose response curve for the *snf1* mutant after 1 hour exposure to TZ at the indicated concentrations and representative images of *snf1* mutant yeast cells colored to reflect the QUEEN ratio where blue/green values are assigned to lower QUEEN ratios (right). Horizontal lines represent the median value. All comparisons between TZ treated samples and the 0 dose were significant at p<0.02. D-E-F) The effect of TZ on ATP and puncta formation in cells expressing Aβ42-GFP. Note that the dose response curve for this batch of TZ batch showed a clear elevation in QUEEN signal at 40 μM. D) Data points represent soluble Aβ42-GFP fluorescence intensity of individual cells lacking puncta after a 4-hour CuSO_4_ induction and a 1-hour treatment with 40 µM TZ; n>800 cells per treatment category; soluble Aβ42-GFP increases with TZ in *snf1* mutants. Horizontal lines depict quartiles. E) Representative images from D are shown with the percentage of cells exhibiting Aβ42-GFP puncta indicated. F) 40 µM TZ increases ATP levels in the *snf1* mutant expressing Aβ42-GFP, based on luciferase-based biochemical measurement of cellular ATP. Significance for all panels was determined by ANOVA analysis with a post-hoc Tukey test (****, p<0.0001; ns, non-significant).

Treatment of the *snf1* mutant with TZ at concentrations between 40–60 μM (the precise value was batch dependent) resulted in a rapid increase in ATP levels (Fig 1C) (Cai et al., 2019a; Schultz et al., 2022, 2024). Treatment of *snf1* mutant yeast with TZ at concentrations that were lower or higher than 40–60 μM were less effective in increasing ATP levels, as expected, given the competition between TZ binding and ATP/ADP binding to PGK1 (Cai et al., 2019a; Riley et al., 2024) (Fig 1C). Of note, WT strains of yeast homeostatically regulate ATP levels and consequently ATP levels were not increased by TZ treatment, although WT strains were included as negative controls in these studies. Interestingly, at TZ concentrations above 50 μM, WT strains showed a slight but significant reduction in ATP levels (Fig S1A). Treatment of *snf1* mutant yeast with tamsulosin, an α_1_-adrenergic receptor antagonist that does not bind PGK1, caused no consistent dose-dependent effect on ATP levels (Fig S1B). These data are in line with past work in cell lines, human induced pluripotent stem cells (IPSC), flies, rodent brain, and *in vivo* in humans with and without PD (Cai et al., 2019b; Schultz et al., 2022, 2024).

Next, we examined the effect of elevated ATP on the aggregation of Aβ42 in the *snf1* mutant expressing Aβ42-GFP under the control of the yeast *CUP1* promoter (Wurth et al., 2002), which can be induced by the addition of copper sulfate (CuSO_4_). This Aβ42-GFP construct is designed to give rise to a diffuse cytoplasmic signal when soluble that is reduced under conditions that promote Aβ42 aggregation (Wurth et al., 2002). First, we confirmed that neither the introduction of Aβ42-GFP expression plasmid nor the induction by CuSO_4_ affected the reduced ATP level that we observed in *snf1* mutant yeast (Fig S1C). Since 40 μM TZ caused a significant elevation in ATP (Fig. 1F) in this instance, we next measured the impact of 40 μM TZ treatment on soluble Aβ42-GFP fluorescence in the *snf1* mutant following CuSO_4_ induction. The *snf1* mutant exhibited a very low level of soluble Aβ42-GFP fluorescence (indicating a high level of Aβ42 aggregation), which was significantly increased by TZ treatment (Fig 1D). In addition, a substantial number of cells in the *snf1* mutant sample (29.5%) contained fluorescent puncta due to Aβ42-GFP aggregation, and the percentage of cells with fluorescent puncta was reduced following TZ treatment (12.6%) (Fig 1E). The increase in soluble signal and reduction in cells with puncta following TZ treatment suggest that the reduced Aβ42-GFP aggregation may be associated with the increased levels of ATP. Of note, studies with the same design were performed in WT yeast strains, and neither the soluble Aβ42-GFP fluorescence signals nor the ATP levels were affected by TZ treatment (Fig 1D, 1F). Critically, TZ treatment did not affect Aβ42-GFP expression levels, as shown by western blot analysis (Fig S1D).

In summary, TZ increased ATP levels in the yeast *snf1* mutant and TZ-mediated elevation in ATP may solubilize Aβ42 aggregates in the *snf1* mutant. The use of yeast mutants to study cellular physiology coupled with the opportunity to conduct simultaneous analyses of ATP levels and protein aggregation on a per-cell basis provide useful supplementary approaches for the investigation of the impact of changing metabolism on aging and disease.

### TZ improves neuropathology and cognitive function in the 5xFAD mouse model of AD

We tested the hypothesis that enhancing glycolysis is neuroprotective against AD in the 5xFAD mouse model of AD. We selected the 5xFAD model because it contains five human AD mutations (three in the Aβ precursor protein (APP) gene, and two in the presenilin gene), and it has reliable amyloid aggregation at ∼2 months of age and deficits in cognitive function assays by ∼5 months. We started TZ treatment (1.2 mg/kg/day in drinking water) in 5xFAD mice at ∼5 weeks of age. Our study also included 5xFAD mice given water without TZ, non-transgenic WT littermate controls treated with TZ, and non-transgenic WT littermate controls given water without TZ. All mice were treated for ∼6 months prior to performing behavioral testing (Fig 2A). All 5xFAD mice treated with TZ survived to 11 months, when they were perfused for histology. Three 5xFAD mice (2 males, 1 female) receiving water died before 11 months; 1 WT male mouse receiving water and 1 WT female mouse receiving TZ died before 11 months. A Cox proportional model revealed no reliable differences in survival between 5xFAD mice receiving TZ or water (p = 0.52; Fig 2B).

**Figure 2:**
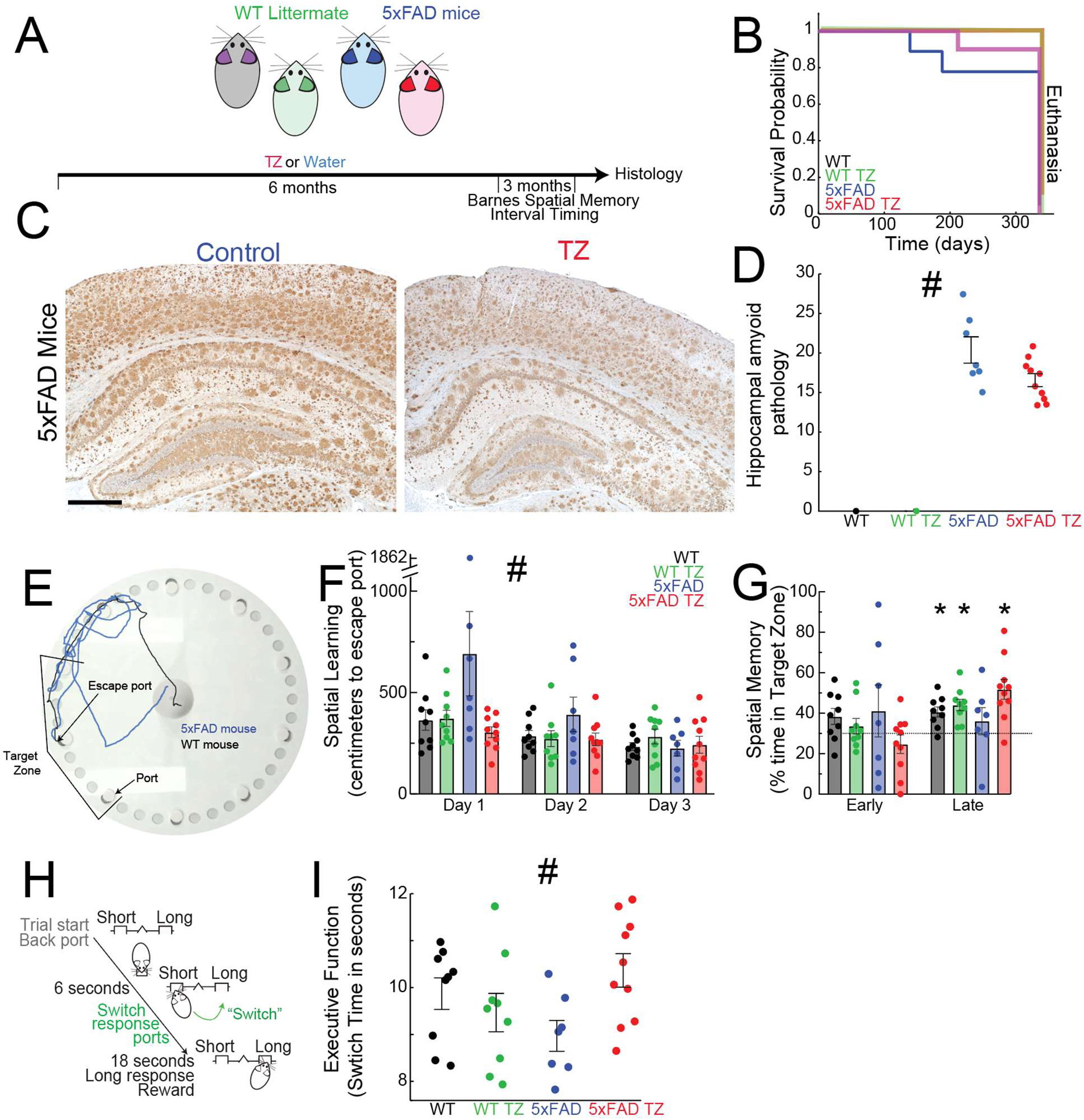
Terazosin (TZ) alleviates amyloid pathology and cognitive deficits in 5xFAD mice. (A) 5xFAD mice and littermate wild-type (WT) controls were treated with TZ or water for 9 months with behavioral assays at 6. (B) Survival of 5xFAD mice vs. WT littermates treated with TZ vs. water. (C) We measured amyloid pathology in the hippocampal CA1 region using 6e10 antibody staining. (D) Amyloid pathology revealed that TZ decreased the amyloid burden in 5xFAD mice. Data from 17 5xFAD mice (10 receiving TZ and 7 receiving water) and 2 control mice (1 receiving TZ and 1 receiving water). (E) We measured spatial learning and memory using the Barnes maze assay in which mice learn which port allows them to escape from an elevated illuminated surface. During testing trials, the distance travelled to the escape port is a measure of spatial learning. A single trial from a 5xFAD mouse (blue) and WT mouse (black) are shown. (F) On Days 1-2, the distance travelled to the escape port was decreased in 5xFAD mice treated with TZ, suggesting that TZ improved the learning of this task. (G) We measured spatial memory by measuring time in the target zone (an area comprising the target and two adjacent ports) on probe trials when the escape port was blocked early in training (on Day 2), and late in training (Day 3). Late in training, 5xFAD mice displayed poor spatial memory with less than >30% in the target zone; this was rescued by TZ. (H) We tested executive function using the switch interval timing task, in which mice estimate an interval of several seconds with a motor response (switch to another port) based on working memory for temporal rules as well as attention to the passage of time. (I) Switch responses, which measure executive function, were early in 5xFAD mice and rescued by TZ. Standard errors are shown on all plots; *p<0.05 via paired t-tests; #p<0.05 from an interaction between 5xFAD and TZ in linear mixed-effects models.

In 5xFAD mice treated with TZ, we examined Aβ pathology through immunohistochemistry using an established amyloid scoring scale, quantified by a veterinary pathologist who was blinded to our TZ treatment system (Maeda et al., 2007; Meyerholz & Beck, 2018a). The amyloid score in the CA1 region of the hippocampus was significantly lower in 5xFAD transgenic mice treated with TZ vs. water (main effect of TZ in 5xFAD mice; p = 0.04; Fig 2C–D). No differences in amyloid scores were observed in the cortex.

We investigated whether TZ improved cognitive performance in 5xFAD mice. First, we performed the Barnes Maze assay of spatial learning and memory (Bertolli et al., 2024; Gawel et al., 2019; O’Leary & Brown, 2013; Weber et al., 2024). The Barnes Maze takes advantage of rodent’s avoidance of illuminated environments and spatial learning and memory for escape routes from an elevated platform. On each trial, mice are placed on an elevated circular illuminated table with 10 ports arranged along the outer edge, one of which allows the mouse to escape to a preferred dark location—the target escape port. As a measure of spatial learning, we quantified how far mice traveled from the start at the center of the illuminated table to the target escape port, averaged across 5 trials on each of 3 days. A shorter distance travelled is associated with faster spatial learning for finding the target (Fig 2E). While all animals decreased the distance travelled over 3 days, 5xFAD mice consistently travelled more distance to the escape port, reflecting slower learning rates. TZ compensated for spatial learning deficits in 5xFAD mice (no main effect of 5xFAD mice; main effect of TZ in 5xFAD mice: p = 0.04; interaction between 5xFAD mice and TZ across all days: p = 0.04; Fig 2F).

We also quantified spatial memory via performance on probe trials, during which the target escape port is blocked. Early in training (24 hours after 1 day of Barnes Maze performance), there was no difference between the percentage of time that all mice spent in the target zone over the blocked escape port (which includes the escape port and two flanking ports; 3/10 ports, or 30% is chance; Fig 2G) implying that Barnes Maze training was insufficient to consolidate spatial memory. However, later in training 24 hours after Day 2 of Barnes Maze performance, WT mice treated with water and with TZ spent significantly more time in the target sector compared to chance (p = 0.03 and p = 0.02, respectively). 5xFAD mice exhibited impaired spatial memory, performing no better than chance (p = 0.66) reflecting impaired spatial memory; however, 5xFAD mice treated with TZ spent significantly more time in the target sector compared to chance (p = 0.004). TZ specifically improved spatial learning and memory, while other behavioral differences between genotypes were unaffected; 5xFAD mice treated with TZ or water travelled less distance overall than WT animals on probe trials (Fig S2). These data indicate that TZ was associated with improved spatial learning and memory in 5xFAD mice.

Finally, to investigate the effect of TZ on executive function, we used an interval timing assay that required mice to estimate an interval of several seconds by making a motor response— switching one nosepoke response port to another. Timing temporal intervals requires working memory for temporal rules as well as attention to the passage of time (Brown, 2006; Parker et al., 2013). In this mouse-optimized version, the timing of when mice switch from one nosepoke response port to the other reflects internal estimates of time because no cue indicates when animals must switch nosepoke response ports (Balci et al., 2008a; Bruce et al., 2025; Weber et al., 2023). Interval timing is impaired in humans with AD as well as 5xFAD mice, which tend to have anticipatory switch times, or earlier switch times, compare to control mice (Carmen Carrasco, 2000; Caselli et al., 2009; El Haj & Kapogiannis, 2016; Gür et al., 2019). Strikingly, TZ compensated for 5xFAD deficits in interval timing (interaction between 5xFAD mice and TZ on switch time: p = 0.02; Fig 2I). Taken together, these data indicate that TZ is protective in animal AD models.

### TZ is associated with a slower decline of cognitive function and FDG-PET biomarkers

To examine if these changes in mice were linked with human AD, we analyzed the Alzheimer’s Disease Neuroimaging Initiative (ADNI), a real-world longitudinal, multi-center, observational study designed to track the progression of AD. ADNI collected sequential neuroimaging and clinical data from a cohort of patients with AD (Jack Jr. et al., 2008). Our preclinical data suggest that TZ may be neuroprotective in AD, and the ADNI database provides a unique resource to test this hypothesis in humans. We found 61 ADNI participants who were taking either TZ or two related drugs that also enhance PGK1 activity (doxazosin (DZ) or alfuzosin (AZ)) and 92 participants who were taking tamsulosin, an α_1_-adrenergic receptor antagonist that does not bind to PGK1. 153 participants taking either TZ, DZ, AZ (TZ/DZ/AZ) or tamsulosin were matched to 306 participants who were not using any of these drugs (non-users; controls). We compared the TZ/DZ/AZ users to the tamsulosin users and to the non-user controls.

We found that the mean annualized rate of change of cognitive function as measured by the Alzheimer’s Disease Assessment Scale-Cognitive 13 (ADAS-CoG 13; a measure of cognitive impairment with higher scores indexing more advanced cognitive impairments) was 1.71 points per year (ppy) (95% CI [1.48 – 1.93]) in patients taking neither drug, which was not significantly different from patients taking tamsulosin (1.62 ppy (95% CI [1.16 – 2.08]; p = 0.74)). Remarkably, the TZ/DZ/AZ group had a slower mean annualized rate of change of the ADAS-CoG 13 (0.96 ppy, 95% CI [0.42 – 1.51]; p = 0.01) compared to the non-user group (Fig 3A). The rate of progression of the ADAS-CoG 13 was also slower in the TZ/DZ/AZ group relative to the tamsulosin group, but the results did not reach statistical significance (p = 0.07). Among all participants, there were no significant group differences in the mean annualized rate of change observed using the Clinical Dementia Rating Scale Sum of Boxes ((CDR-SOB), which measures the severity of dementia (non-users control group: 0.62 ppy (95% CI [0.55 – 0.68]); TZ/DZ/AZ group: 0.53 ppy (95% CI [0.37 – 0.69]); tamsulosin group: 0.56 ppy (95% CI [0.43 – 0.69]). These data suggest that TZ/DZ/AZ could mitigate some aspects of cognitive impairment in patients with AD.

**Figure 3.**
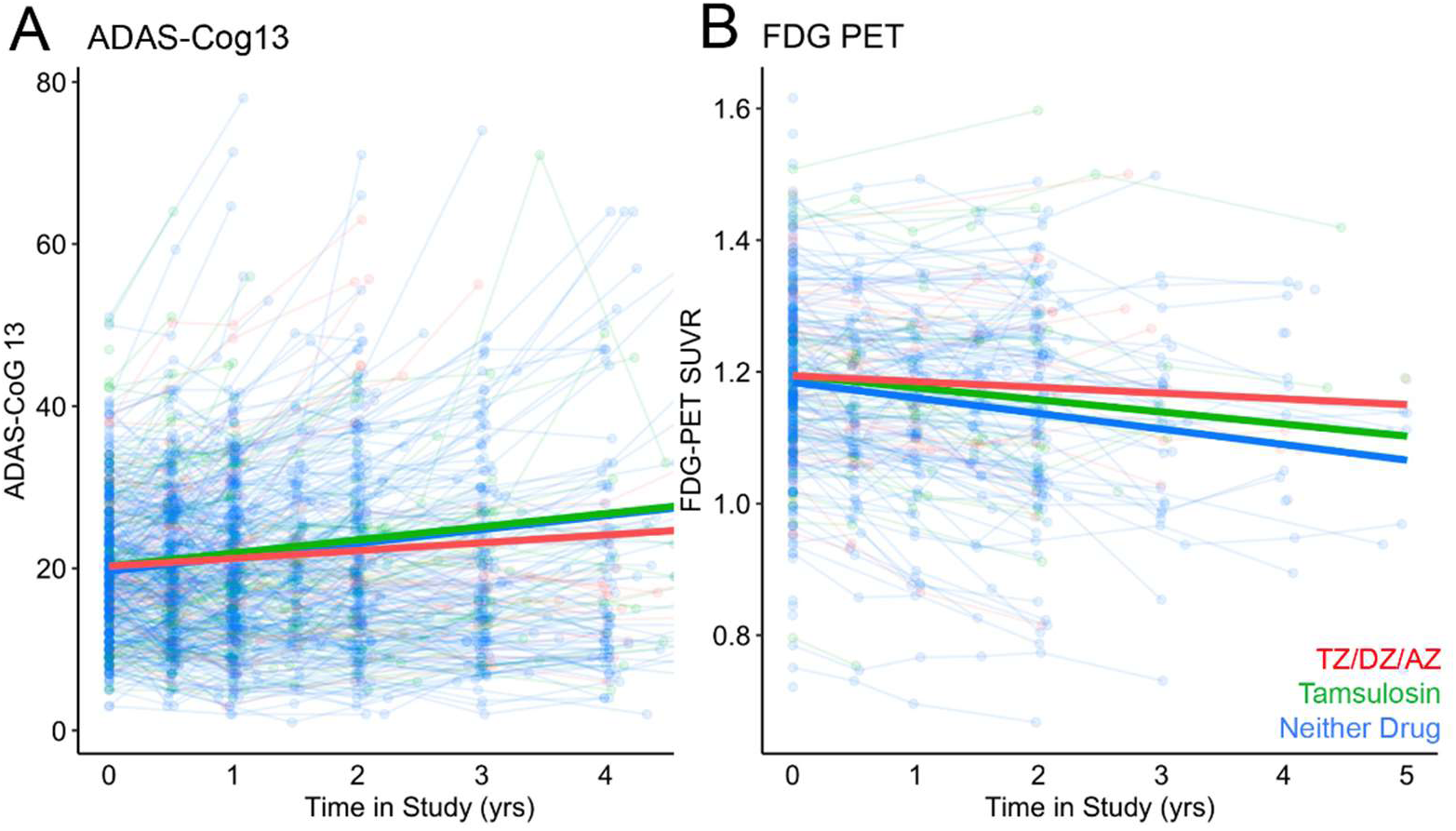
Patients with Alzheimer’s disease taking TZ/DZ/AZ had slower cognitive decline and slower progression of ^18^F-FDG-PET abnormality. We analyzed the Alzheimer’s Disease Neuroimaging Initiative (ADNI) database and found slower cognitive decline (A), as indicated by an increase in the ADAS-Cog 13 score, and less decrease in 18F-FDG-PET (B) in AD patients taking TZ/DZ/AZ compared with AD patients taking tamsulosin or none of these drugs; note that fewer patients had PET scans later in the study.

The ADNI database also includes extensive neuroimaging biomarkers, including a measure of brain metabolism, ^18^F-FDG-PET (Mosconi, 2005a). In line with a slower progression of cognitive impairment, we found that the rate of decline in the FDG-PET signal was significantly slower in the TZ/DZ/AZ group (mean annualized change = -0.009, 95% CI [-0.019 – 0.002]; (p = 0.012)) compared to non-user controls (mean annualized change = -0.02, 95% CI [-0.028 – -0.019]; Fig 3B). The tamsulosin group did not reliably differ compared to the other two groups (mean annualized change = -0.02, 95% CI [-0.029 – -0.008]). These data provide additional evidence that glycolysis-enhancing drugs are protective in AD.

### TZ protects against the development of AD

We investigated a large-scale administrative insurance claims database to explore whether TZ is associated with a lower risk of AD. Using the Merative Marketscan Commercial Claims and Encounters and Medicare Coordination of Benefits databases of insurance claims, we identified 1,028,561 users of TZ, DZ, or AZ (TZ/DZ/AZ); 2,854,989 users of tamsulosin; and 653,508 users of 5-α-reductase inhibitors (5ARI), a drug also used in men with benign prostatic hyperplasia (BPH). We used an active-comparator and new-user design, the gold standard in observational pharmacoepidemiological studies (Lund et al., 2015), which has not been reliably used in retrospective studies of BPH drugs and dementia (Duan et al., 2018; Fung et al., 2024; Sasane et al., 2021). After applying our exclusion criteria, we had a final cohort of 209,146 new users of TZ/DZ/AZ, 1,013,108 new users of tamsulosin, and 120,431 new users of the 5ARIs finasteride or dutasteride (Table S1).

New users of TZ/DZ/AZ were followed for 594,862 person-years, and we observed 7,054 new cases of AD (11.9 cases per 1,000 people per year). This rate was lower than the rates observed among new users of tamsulosin (2,761,034 person-years; 46,915 cases; 17.0 cases per 1,000 people per year) or 5ARI (352,377 person-years; 4,586 cases; 13.0 cases per 1,000 people per year). Correspondingly, there was greater AD-free survival among TZ/DZ/AZ users compared to either tamsulosin or 5ARI users (Fig 4). The estimated hazard ratio for AD for TZ/DZ/AZ users was lower compared to tamsulosin users (HR = 0.69, t = -29.4, p < 10^−16^, 95% CI: [0.67 – 0.71]). Critically, the hazard of AD was also lower for TZ/DZ/AZ users compared to 5ARI users (HR = 0.77, t = -18.2, p < 10^−16^, 95% CI: [0.74 – 0.88]) demonstrating that these results could not be explained by effects of tamsulosin alone (Simmering et al., 2022).

**Figure 4.**
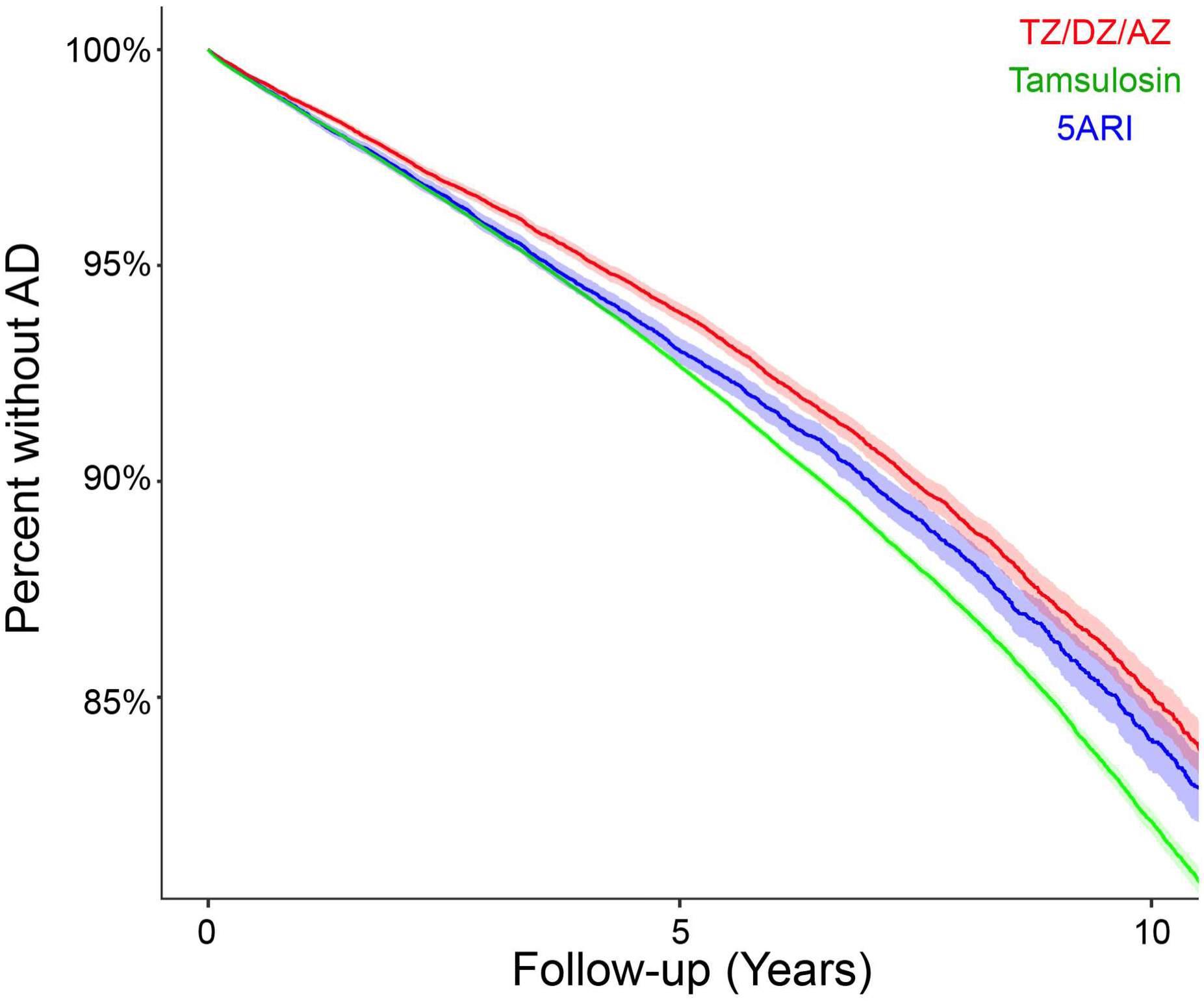
Patients taking TZ/DZ/AZ have less risk of Alzheimer’s disease (AD) in the Merative Marketscan human database of insurance claims. A Kaplan-Meier curve of the probability of AD diagnoses calculated from the Merative Marketscan dataset for users of TZ/DZ/AZ (red) vs. tamulosin (green) or 5ARI (blue). Patients using TZ/DZ/AZ were *less likely* to get AD than patients using tamsulosin or 5ARI.

There were some notable differences between the groups at baseline (Table S2). For instance, users of TZ/DZ/AZ were much more likely to have hypertension, renal disease, and diabetes with complications. Additionally, TZ/DZ/AZ users were likely to be older (average age of 62.0 years for TZ/DZ/AZ users vs. 60.2 years for tamsulosin users [standard mean difference (SMB) = 0.16] and 61.5 years for 5ARI users). [SMD = 0.04]). Meanwhile, 5ARI users were much more likely to have abnormal prostate-specific antigen (PSA) values (SMD for TZ/DZ/AZ versus 5ARI = 0.16, tamsulosin versus 5ARI = 0.14) and a benign prostatic hyperplasia (BPH) diagnosis (SMD for TZ/DZ/AZ versus 5ARI = 0.09, tamsulosin versus 5ARI = 0.09). Users of tamsulosin had an average of 2.15 diagnoses per health care encounter, more than either TZ/DZ/AZ users (1.82, SMD = 0.37) or 5ARI users (1.81), SMD = 0.39). These differences were mitigated through our weighting algorithm and propensity-score matching (Table S3). Due to the entropy balancing approach used, the means were exactly equal and the range of the weights were narrow. All confounders were balanced with similar variabilities. The lookback period and follow-up durations, which were not included in the inverse probability of treatment weighting (IPTW) adjustment, were similar across all three groups.

The association between a lower hazard of AD and the use of TZ/DZ/AZ persisted after adjustment. In a weighted pseudo-population, we observed 7,200 cases of AD in 533,753 person-years of follow-up among new users of TZ/DZ/AZ (13.5 per 1,000 people per year). Users of 5ARI (322,522 person-years, 4,887 cases, 15.2 cases per 1,000 people per year) and users of tamsulosin (2,870,393 person-years, 47,228 cases, 16.5 cases per 1,000 people per year) had higher incidence rates. The survival curve (Fig 4) shows lower AD incidence after adjustment through IPTW. The Cox proportional hazards model shows a lower hazard of AD in new users of TZ/DZ/AZ compared to new users of tamsulosin (HR = 0.84, t = -867.9, p < 10^−16^, 95% CI: [0.84 – 0.84]) and a smaller difference between users of 5ARI and tamsulosin (HR = 0.93, t = - 385.5, p < 10^−16^, 95% CI: [0.93 – 0.93]).

Inspection of the Schoenfeld residuals indicated violations of the proportional hazards assumption. We re-estimated our model with time-varying coefficients (Table S4). Across all time intervals, compared to users of tamsulosin, users of TZ/DZ/AZ had a reduced hazard of AD (HR of 0.80–0.85) that was consistent with the overall HR estimate (0.84). On the other hand, the relative hazard between users of 5ARI and tamsulosin varied over follow-up. During the first year of follow-up, there were no reliable difference between users of 5ARI and tamsulosin (HR = 0.98); however, as follow-up time increased the hazard among the users of 5ARI decreased (e.g., HR = 0.86 for 7.5 to 10 years). After inclusion of the time-varying coefficients, the Schoenfeld residuals indicated no violations of the proportional hazards assumption (p = 0.61).

Finally, we performed additional sensitivity analyses (Table S5). The estimated associations were effectively unchanged when patients with a neurological history were excluded, or a 1-year delay in follow-up was implemented. An enhanced case definition requiring two or more AD diagnoses persisting for at least 1 year yielded a greater reduction in AD diagnosis for users of TZ/DZ/AZ compared to our main model, which used any diagnosis of AD. As expected, incidence rates of AD were six-fold higher among people with a neurological history; however, the reduction in AD hazard associated with the use of TZ/DZ/AZ was considerably smaller in the model requiring no prior neurological history, compared to the main model of any diagnosis of AD. Taken together, these data demonstrate that TZ/DZ/AZ may be protective against the development of AD when compared to both tamsulosin and 5ARI and imply that TZ/DZ/AZ may be neuroprotective in AD.

## DISCUSSION

We tested the hypothesis that enhancing glycolysis is neuroprotective in AD. We found that the glycolysis-enhancing drug TZ 1) increased ATP and decreased Aβ aggregation in yeast, 2) decreased hippocampal Aβ pathology in 5xFAD mice and protected behavioral performance of 5xFAD mice in the Barnes Maze assay of spatial memory and in the switch interval timing assay, 3) decreased rates of cognitive decline and slowed the progression of FDG-PET imaging abnormality of participants in the ADNI dataset, and 4) decreased the risk of developing AD in the Merative Marketscan human dataset of insurance claims. These data provide convergent evidence that enhancing glycolysis with TZ has the potential to be neuroprotective in AD. While testing the efficacy of glycolysis-enhancing drugs such as TZ will require rigorous clinical trials, our data implicate a novel mechanism of neuroprotection that is relevant for AD.

AD presents a significant public health challenge, with energy metabolism impairment recognized as a key pathological feature contributing to its progression (Mosconi, 2005a, 2005b; Profenno et al., 2010). In recent years, the exploration of TZ, primarily used as an α_1_-adrenergic receptor antagonist in hypertension and BPH, has revealed neuroprotective properties beyond its traditional indications. TZ binds to PGK1 and increases ATP in cell lines, in flies, and in preclinical mouse models (Cai et al., 2019b; X. Chen et al., 2015; Kokotos et al., n.d.; Weber et al., 2024). TZ augments cellular energy metabolism, which may be crucial in mitigating neurodegenerative processes through a variety of mechanisms (Cai et al., 2019a; Chaytow et al., 2022; H. Chen et al., 2023; Schultz et al., 2022; Simmering et al., 2021, 2022; Yu et al., 2022). Our ability to study these relationship in yeast is exciting because it might identify fundamental principles of how ATP interacts with protein aggregation, which will lead to a further understanding of pathological mechanisms and interventions (Patel et al., 2017; Takaine, 2019b; Takaine et al., 2022).

This work is supported by a recent report demonstrating that TZ was neuroprotective in another AD mouse model (Yu et al., 2022). This group also found similar effects while manipulating the α_1_-adrenergic receptor and interpreted their data in this context. Our past work has shown that TZ interacts directly with PGK1, independent of its α1-adrenergic receptor effects, and that other drugs with α1-adrenergic receptor antagonism do not interact with PGK1, do not increase cellular ATP, and are not neuroprotective (Cai et al., 2019b; X. Chen et al., 2015; Weber et al., 2023).

Our data extend to real-world human observational data from ADNI and Merative to investigate the clinical relevance of TZ in AD progression. The ADNI dataset provided valuable insights into the longitudinal effects of TZ on the progression of AD. Participants with AD who were treated with TZ/DZ/AZ exhibited a slower progression of cognitive dysfunction measured by ADAS-Cog 13 scores and a slower decline in FDG-PET signal, indicative of preserved neuronal metabolic activity. These findings suggest that TZ may not only slow cognitive decline but also preserve functional brain activity associated with AD pathology. Analyses of the Merative Marketscan database revealed a lower hazard of developing AD among patients prescribed either TZ or a related glycolysis-enhancing drug (AZ or DZ) compared to those prescribed tamsulosin, an α1-adrenergic receptor antagonist and 5ARI without glycolysis-enhancing properties. This retrospective cohort analysis underscores the potential clinical benefit of glycolysis-enhancing drugs in reducing AD risk and highlights TZ as a promising candidate. Of note, a recent study found that tamsulosin increased the risk of dementia (Duan et al., 2018). However, this result has been critiqued for the following factors: 1) the diagnosis of dementia was not specific to AD, 2) tamsulosin does not cross the blood brain barrier, 3) prescription habits varied between comparison groups, and 4) an inclusion of an active comparator is the gold standard design for validity (Andrade, 2018; Tae et al., 2019; Ulrich, 2018). In our recent work, we found that TZ/DZ/AZ were protective in PD when compared to multiple comparators, including tamsulosin and 5ARIs (Simmering et al., 2022). This inclusion of active comparisons is critical in our analysis, as patients not taking any drugs for BPH are categorically different from patients who take drugs for BPH (suggesting that they have routine access to healthcare). Our current analysis of TZ/DZ/AZ in AD accounts for these factors (Simmering et al., 2021b, 2022).

In the Merative Marketscan human database, our results found an association between the use of TZ/DZ/AZ and a lower risk of AD compared to the use of either tamsulosin or 5ARIs. This effect persisted and was largely unchanged in several sensitivity analyses using alternative case definitions, delayed follow-up starts, or important sub-groups of interest. The effect of the use of TZ/DZ/AZ is temporally consistent with time-varying coefficients, showing a hazard ratio between 0.80 and 0.85 for 10 years after the medication start date, compared to that seen with tamsulosin use. Interestingly, in two of the sensitivity analyses, the results pointed toward a plausible causal interpretation of these results. First, we observed a slightly lower hazard of AD diagnosis for users of 5ARI compared to tamsulosin. However, it became clear that the association between 5ARI use and AD hazard was time-varying (Table S4). The time-varying coefficients show no difference in the hazard of AD between users of 5ARI and users of tamsulosin in the first 3 years following the drug start date. It seems unlikely that a causal effect of 5ARI on AD risk would only manifest 3 years after treatment start. However, use of TZ/DZ/AZ is consistently associated with a lower hazard of AD at all time points. This supports an interpretation that 5ARI and tamsulosin are true “control” treatments with no effect on AD risk. Second, we found a muted effect of TZ/DZ/AZ in a subgroup with neurological comorbidities, which could include patients on a path toward an AD diagnosis (Table S5). The incidence rate of AD in this subgroup of patients is much higher than that of people without neurological comorbidities (∼65 per 1,000 people per year compared to ∼13 per 1,000 people in our data). In this subgroup of patients, TZ/DZ/AZ is still associated with a lower hazard of AD, but the effect is smaller (compared to tamsulosin users, HR = 0.90 vs. 0.84 in the overall population). It is likely that this population is further along the pathogenesis of AD, and we would expect any neuroprotective effect of PGK1 activation to be correspondingly smaller. Slowing the progression of AD in a person on the threshold of an AD diagnosis is unlikely to result in as much of a reduction in the hazard of AD as it would in a person who is still very early in the prodromal stage toward the development of AD.

Our results have several limitations. First, there are many mouse models that capture aspects of human AD, and our work supports prior findings in the APP/Swe model (Yu et al., 2022). Second, we report that in the ADNI database, compared to tamsulosin, TZ/DZ/AZ nonsignificantly improved the CDR-SOB and slowed AD progression as measured by FDG-PET, although these differences were not statistically significant. It is compelling that TZ/DZ/AZ is distinct from controls in the ADAS-Cog13 and FDG-PET, and that it provides a hypothesis-driven signal in a well-described AD cohort. Finally, the retrospective nature of our Merative Marketscan database analyses and inherent biases in ADNI may influence the interpretation of our results. We used a rigorous analytical framework with the active-comparator and new-user design to further reduce confounding by IPTW and included two active comparators. However, our ability to control confounding and obtain valid estimates depends on the accuracy and completeness of data observed. Providers may have important considerations when prescribing TZ/AZ/DZ vs. tamsulosin, such as hypotension or general frailty, that are correlated with AD risk but are not recorded in the claims record. Indeed, despite these drugs being primarily used for the treatment of BPH, most users are never diagnosed with BPH. It is possible that important confounders are omitted, incompletely recorded, or otherwise unknown. Although we included alternative case definitions, delayed follow-up, and subgroup analyses, and we observed consistent results across all these analyses, the risk of omitted confounders remains. We do not have detailed clinical data on these patients included in our analyses, such as MoCA scores, which may be a more sensitive measure of neuroprotection and progression. While some of these concerns might be addressed with more advanced retrospective analyses, they can only definitively be addressed with a prospective clinical trial, although the numbers and design of this trial would be necessarily large and costly.

In summary, we provide evidence that glycolysis-enhancing drugs increase ATP and decreased amyloid aggregation in yeast mutants. Furthermore, we find that glycolysis-enhancing drugs mitigate amyloid aggregation and preserve cognitive behaviors in the 5xFAD mouse model of AD. Furthermore, in ADNI we find slower progression of the ADAS-Cog13 and FDG-PET in patients taking glycolysis-enhancing drugs, and in the Merative Marketscan retrospective human dataset, we find that patients taking these drugs have decreased risk of being diagnosed with AD. Our results suggest that enhancing glycolysis is neuroprotective in AD and could inspire future clinical trials of repurposed glycolysis-enhancing drugs and new research into novel AD-relevant mechanisms.

## MATERIALS AND METHODS

### Yeast

#### Strains

WT and *snf1* mutant strains with a 3x integrated QUEEN reporter (MTY3264 and MTY3371 (Takaine et al., 2022) were purchased from the Yeast Genetic Resource Centre Japan (YGRC). QUEEN reporter strains were grown to mid-log phase from fresh overnight cultures in synthetic complete supplement mixture (SC) media at 25 °C. The *snf1* mutant strain and the corresponding WT strain (BY4730) that were transformed with the Aβ42-GFP reporter (see plasmids below) were from the yeast deletion collection (Brachmann et al., 1998). The transformed strains were grown to mid-log phase from fresh overnight cultures in SC-URA (SC media lacking uracil) media at 30 °C. Terazosin (TZ) HCl dihydrate was purchased from Selleck Biochemicals (S2059). Stocks were prepared at 20 mM in water and stored as aliquots at -80 °C. TZ was added to log phase cultures at final concentrations of 0–300 µM for dose response curves, with ATP being evaluated 60 minutes after each TZ addition. TZ was added at a final concentration of 40 µM to Aβ42-GFP reporter strains following 3 hours of reporter induction with CuSO_4_. Sixty minutes after TZ addition, GFP fluorescence, biochemical ATP values, and protein expression were evaluated using western blot analysis.

#### Yeast plasmids

The Aβ42-GFP yeast expression plasmid was constructed by PCR amplification of the Aβ42-GFP cassette from plasmid aβ-GFP (pET28(A+)) (Addgene 72203) (Wurth et al., 2002). The PCR product was co-transformed into yeast along with a linearized yeast P_CUP1_ vector. The resulting recombinant construct, pGP2320, was isolated from yeast, confirmed by sequencing, and introduced into the test wild type and *snf1*Δ strains indicated in Fig. 1.

#### Yeast microscopy and QUEEN ratiometric analysis

Imaging was performed at 25 °C using fluorescence microscopy. Bright field, differential interference contrast (DIC), and two fluorescence images (Fluorescein (FITC) (ex480 image) and Ex409 (ex409 image)) were taken from multiple fields. The selection of fields was conducted under DIC to avoid unconscious bias. Open-source Fiji software was used to calculate the QUEEN ratio (Takaine et al., 2019).

To measure GFP fluorescence of Aβ42-GFP, cells were circled in the DIC channel and ROIs were measured in GFP for raw integrated density (“RawIntDen”) and area of the shape (“Area”). Similarly, five areas of the image background were measured to correct for background fluorescence in reported data points. The raw integrated density was divided by the area to establish fluorescence intensity for both cells and background in each field. The average background fluorescence intensity was subtracted from this value for each cell reported to provide the corrected fluorescence intensity (A.U./mm^2^) reported in Fig 1D. Fluorescence values are reported for cells having only soluble signal (i.e., no punctate GFP).

#### Yeast biochemical ATP measurements

Measurements of ATP in yeast cell extracts were performed as in (Xu & Bretscher, 2014)with minor modifications using the Invitrogen ATP Determination Kit (Invitrogen, Carlsbad, CA, #A22066). Briefly, extracts were prepared from late-log phase cultures (OD_600_ = 0.9–1.0), and the number of cells was quantified by hemocytometer. ATP was extracted by treating the pellets with a 10% trichloroacetic acid (TCA) solution. The cell suspension was vortexed for 1 minute and incubated on ice for 10 minutes. The pH of the suspension was adjusted to 7.2 by addition of 25 mM Tris/2 mM EDTA pH 9.4 to prevent inhibition of the luciferase reaction. Then 10 μl of extract was measured in 90 μl of the Invitrogen kit assay, and the ATP concentration was determined using an ATP standard curve. Luciferase was measured using a BioTek Cytation5 (Agilent, Santa Clara, CA) cell imaging multimode reader.

#### Yeast extracts and western blot analysis

Extracts from Aβ42-GFP-expressing yeast were prepared from log-phase cultures using a glass bead lysis method. Cultures used for imaging were then pelleted and frozen at -80 °C and used within 1 week. Approximately 100 µl of glass beads and 100 µl of extraction buffer (30 mM Tris, pH 8.5, 5 mM EDTA, 3 mM DTT, 5% glycerol plus 2.5 µl of 40 mM PMSF, and 2.5 µl of PIC; Sigma, P8215) were added to thawed pellets. Cells were disrupted with 3 × 60 seconds of Qiagen Tissue Lyser LT (Hilden, Germany) cycles alternating with 1 minute on ice. The mixture was centrifuged at 13,000 rpm for 1 minute, and the beads and pellet were discarded. The protein concentration was determined using the Bradford Protein Assay (Bio-Rad, Hercules, CA) with a bovine serum albumin (BSA) standard curve. Each extract was mixed with 5X LSB (Lauryl sulfate broth; 15% SDS, 0.575 M sucrose, 0.325 M Tris-HCl, pH 6.8, 5% β-mercaptoethanol, and 0.002% bromophenol blue) and incubated at room temperature (RT) for 15 minutes before being loaded on gradient Tris-Glycine polyacrylamide gels (Invitrogen) and subject to electrophoresis in Tris-glycine buffer + SDS at 100 volts for 90 minutes. Protein was transferred to nitrocellulose in Tris-glycine buffer (25 mM Tris, 192 mM glycine, 20% methanol) by electroblotting. Post-transfer membranes were incubated with Ponceau S for lane-to-lane normalization. Membranes were washed once in phosphate-buffered saline (PBS)-T (1X PBS with 0.1% Tween-20) to remove the stain prior to blocking. Membranes were blocked with 5% milk in 1X PBST for 1.5 hours at RT with rocking. α-GFP antibody (Proteintech, Rosemont, IL) was used at a 1:10,000 dilution. Secondary antibody (1:20,000) was IRDye 800CW goat anti-rabbit IgG (LI-COR). Membranes were imaged on the LI-COR Odyssey Fc imaging system in the 800 channels (IRDye800) with 2-minute exposure times. The GFP signal was normalized to the Ponceau S intensity on a per-lane basis.

### Mice

#### 5xFAD mice

All experimental procedures were performed in accordance with the relevant guidelines of Protocol #0062039 and with the approval of the Institutional Animal Care and Use Committee (IACUC) at the University of Iowa. 5xFAD transgenic mice (Tg(APPSwFlLon,PSEN1 *M146L*L286V)6799Vas) (Oakley et al., 2006) and non-transgenic littermates were received from The Jackson Laboratory (Bar Harbor, ME) at approximately 5 weeks of age and were communally housed on a 12-hour light/dark cycle with *ad lib* access to laboratory rodent chow and water. To facilitate operant behavioral training (described below), mice were weighed daily and maintained on a restricted diet with *ad lib* access to water.

#### TZ administration

TZ (Tocris #1506, Minneapolis, MN) was delivered to mice through *ad lib* access to treated water in plastic water bottles, starting at 5 weeks of age and continuing until mice were euthanized. The vehicle control group had access to regular drinking water contained in the same type of plastic water bottles. Preparation of TZ began with a 41-mM stock solution (18.8 mg/ml of TZ HCl-dihydrate in water ∼17.3 mg/ml of TZ HCl) stored at 4 °C for up to 1 week. We added 106 μL of stock solution, equating to 1.86 mg TZ HCl, to 300 ml water, to which mice had free access. The concentration of the final drinking water was 6.2 μg/ml TZ (1.9 mg/300 ml). An average mouse drinks ∼5 ml of water per day (Nicolaus et al., 2016), so each mouse consumed ∼31 μg TZ daily (6.2 μg/ml x 5 ml/day). Additionally, experimental mice weigh on average 0.025 kg; thus, the TZ dose received by our animals was ∼1.23 mg/kg/day (31μg/day/0.025 kg). All water bottles containing TZ or water were replaced every 2 days for the duration of the experiment until mice were euthanized (Cai et al., 2019a).

#### Barnes maze spatial learning and memory assay

At 8 months of age, mice were run on the Barnes maze assay in balanced cohorts using a 3-day optimized protocol for procedural learning and spatial memory acquisition (Bertolli et al., 2024). Briefly, during the Barnes Maze assay, a mouse is placed on an elevated circular illuminated table with 10 ports arranged along the outer edge, one of which allows the mouse to escape to a preferred dark location—the target escape port. This protocol uses 3D-printed escape shuttles attached to the target escape port, combined with rotating the entire maze at each trial to reduce the possibility of unwanted cues interfering with spatial memory acquisition. On Day 1, mice performed one unanalyzed habituation trial and five spatial learning trials, with homecage bedding placed in the shuttle box at the bottom of all escape ports (target and non-target). The maze was turned every trial to minimize odor as a navigation cue. Intertrial intervals were 3 minutes from the time animals escaped. On Day 2, mice performed one probe trial to test early spatial retention ∼24 hours after Day 1; on probe trials all ports were blocked to assess whether animals remember the location of the target escape port. Probe trials were followed by five spatial learning trials in which the escape port was unblocked. On Day 3, mice performed one probe trial to test spatial memory retention followed by five spatial learning trials. Data from the five spatial learning trials on each day were analyzed by averaging the distance travelled to reach the escape port. Data from the probe trials on Day 2 (“early”) and Day 3 (“late”) were analyzed as the percentage of time investigating the target spatial location, defined as a pie slice encompassing the target escape port and the two adjacent ports (3/10^th^ of the maze) during a 90-second probe, and compared to a theoretical mean of 30%.

#### Switch interval timing assay

Our mouse-optimized switch interval timing task is designed to capture an animal’s internal representation of time as detailed in our prior work (Balci et al., 2008b; Bruce et al., 2021, 2025; Larson et al., 2022; Tosun et al., 2016; Weber et al., 2023, 2024). Mice are trained to switch from a “short” to a “long” nosepoke after approximately 6 seconds. This switch is an explicit time-based decision that requires working memory for temporal rules and attention to the passage of time; this time-based decision to switch can be employed to model cognitive deficits in neurodegenerative disease (Gür et al., 2020; Larson et al., 2022; Parker et al., 2015; Singh et al., 2021).

Briefly, mice were trained in standard operant chambers enclosed in sound-attenuated cabinets (MedAssociates, St. Albans, VT) that contained two light-equipped nosepoke response ports (left and right) on the front wall, a reward hopper (where food rewards are delivered) located between the two nosepoke response ports, and another light-equipped nosepoke on the back wall, opposite the reward hopper. Operant training began by “shaping” the animal’s behavior for 3 days. Trial initiation began with a response at the back nosepoke, at which point either the left or right front nosepoke was illuminated, and a response at the appropriate port resulted in a reward. After shaping, mice were then advanced to the interval timing switch protocol, for which each session was randomly organized into 50% short (6 seconds) and 50% long (18 seconds) trials. Either the left or right nosepoke was designated for short trials, and the contralateral nosepoke was designated for long trials (counterbalanced across mice and experimental groups). For both trial types, the trial is initiated with a back nosepoke response that generates two identical light cues above both the left and right nosepokes, along with an 8-kHz tone (72 dB). In short trials mice receive a reward after 6 seconds for the first response at the designated nosepoke. In long trials, mice begin by responding at the designated short-trial nosepoke but do not receive a reward. When there is no reinforcement after 6 seconds at the short-trial nosepoke, mice must switch to the designated long-trial nosepoke until a reward is delivered after 18 seconds. Median switch and CV times are plotted in Fig 2I-J. Only long switch trials were analyzed. Behavioral performance was analyzed for the last 4 days of training.

#### Histology

Mice were anesthetized with ketamine (100 mg/kg intraperitoneal (IP)) and xylazine (10 mg/kg IP) and transcardially perfused with cold PBS and 4% paraformaldehyde (PFA). Brains were removed and post-fixed in 4% PFA overnight, followed by immersion in a 30% sucrose in a PBS solution for at least 48 hours before 4–5-µm sections were collected and embedded in paraffin (Larson et al., 2022). The Comparative Pathology Laboratory at the University of Iowa performed all sectioning and immunohistochemistry to visualize amyloid pathology in the mouse brain (Lee et al., 2022). Antigen retrieval was done using citrate buffer (pH 6.0) in the decloaker (Biocare Medical, Pacheco, CA) at 110 °C for 15 minutes, followed by incubation in 6e10 monoclonal antibody (NBP2-62566, Novus, Centennial, CO) (1:5,000 in Dako diluent buffer) for 30 minutes at room temperature, and by incubation with the secondary antibody (anti-rabbit, HRP conjugated + Envision, Agilent) for 30 minutes. Sections were mounted, counterstained with hematoxylin for 3 minutes, and coverslipped. Slides were visualized using an Olympus BX53 microscope, DP73 digital camera, and CellSens Dimension Software (Olympus, Tokyo, Japan).

For amyloid pathology quantification, the pathologist (DM) examined tissues using the post-examination method of masking to group assignments (Meyerholz & Beck, 2018b). To assess in an unbiased approach, the slides were masked and examined for immunostaining using specialized software (Olympus BX53 microscope, DP73 camera, and CellSens Dimension Software). Areas of interest were defined as hippocampus (CA1 region from dentate gyrus to cortex) and cortex (immediately adjacent to the hippocampus region). These defined regions were evaluated with automated thresholding to detect 3,3’-Diaminobenzidine (DAB) staining, and the output was the percentage of the area of tissue that was positive. These percentage data were transformed and normalized relative to staining in the transgenic-vehicle group using the Graphpad Prism Software (Graphpad, San Deigo, CA).

#### Statistics

We analyzed effects of TZ in amyloid pathology and interval timing using linear mixed effects models (Weber et al, 2023). Evaluation of learning rate for Barnes was analyzed with pre-planned comparisons at each day. Probe trials were tested for statistical evidence of learning by the group, by comparing the percent time spent in the target quadrant to a theoretical mean (30%) representing chance performance. (Bertolli et al., 2024). Four sessions of interval timing behavior were collected and analyzed per mouse. All data and statistical approaches were reviewed by the Biostatistics, Epidemiology, and Research Design Core (BERD) at the Institute for Clinical and Translational Science (ICTS) at the University of Iowa. All code and data are made available at: http://narayanan.lab.uiowa.edu/article/datasets. An α of p < 0.05 was considered significant.

#### ADNI database analysis

The Alzheimer’s Disease Neuroimaging Initiative (ADNI) is a multi-site observational study of patients with mild cognitive impairment (MCI) or AD. Data from the ADNI database are made publicly available, and at the time of our analysis, there were 2,343 participants. The ADNI study collects extensive data, including drug use, clinical outcome measures related to AD disease progression, and multiple neuroimaging sequences, among others.

In the current study, we aimed to compare clinical and neuroimaging outcome measures between eligible ADNI participants who were using TZ/DZ/AZ, tamsulosin, or none of these drugs. To be eligible for inclusion in these analyses, participants had to be male (females using TZ/DZ/AZ were less common and more likely to use these drugs for other indications); have more than one eligible ADNI visit; and carry a diagnosis of MCI or AD at their first eligible ADNI visit.

Use of drugs of interest (TZ/DZ/AZ or tamsulosin) was defined as at least 30 days of use during the time that the participant was enrolled in the ADNI dataset. If the participant started and stopped one of the drugs of interest prior to entering the study, they were not considered to be using the drug. Most participants were already taking the drug of interest at the time that they entered the study; however, some began treatment during their participation in the ADNI study. In these instances, the first visit that the participant was known to be taking the drug of interest was considered to be the baseline visit. Concomitant use of TZ/DZ/AZ and tamsulosin was prohibited for these analyses. If a participant switched from one drug class to another, only the time of their first eligible exposure was included, and subsequent visits after switching were discarded.

Given that male participants who would be prescribed TZ/DZ/AZ or tamsulosin were of similar age and had similar comorbidities, all these participants were grouped together. We then performed 2:1 propensity score matching using a nearest neighbor technique to match users of TZ/DZ/AZ or tamsulosin to participants who had never been exposed to either drug group. Matching was based on baseline values of age, ADAS-Cog13 scores, CDR-SOB, baseline diagnosis (MCI or AD), and the number of copies of the APOE4 allele, which is a key modifier of AD recorded in the ADNI database.

Most participants were included in the ADNI study for 5 or fewer years. After matching, we constructed linear mixed effects regression models to compare the trajectory of change of the CDR-SOB, the ADAS-Cog13, and the mean annualized change in FDG-PET standardized uptake value, among participants taking TZ/DZ/AZ or tamsulosin and matched participants who used none of these drugs. The models included a per participant intercept, and the predictor of interest was the group*duration interaction. Specifically, “duration” was the number of years in the ADNI study for each participant.

All analyses were performed in RStudio version 4.2.2. Matching procedures were carried out using the *MatchIt* package (version 4.5.0), and LMER analyses were performed using the *lmerTest* package (version 3.1.3). Results were considered significant if the *p* value was <0.05.

### Merative Marketscan administrative data of insurance claims

#### Cohort construction

We used a new-user, active-comparator design. This is the gold standard design for observational pharmacoepidemiological investigations, as it reduces many important possible confounders, especially confounding by indication (Lund et al., 2015). We identified all dispensing events for TZ, AZ, DZ, tamsulosin, finasteride and dutasteride. We grouped these drugs into three bins: TZ/DZ/AZ, tamsulosin, and finasteride/dutasteride. To ensure that we were identifying new users, we required at least 365 days of enrollment in the Merative Marketscan Commercial Claims and Encounters or Medicare Coordination of Benefits databases with prescription drug coverage before the first dispensing event. The first dispensing event was identified as our “index date,” and our “lookback period” was defined as the period before the index date, during which an individual was enrolled in an insurance plan contributing data to the Merative Marketscan database.

We then applied several exclusion criteria. First, the individual could not switch between the drug groups. Second, we required at least 1 day of follow-up after the index date. Third, these drugs are primarily used to treat BPH, with other indications being less common (TZ is used as a late-stage treatment for hypertension; tamsulosin can be used in the medical management of kidney stones). As a result, we decided to exclude female users. While most male individuals taking these drugs are likely using them for the management of BPH, no female users are managing BPH. Excluding women reduces unobserved heterogeneity, albeit at the cost of some representativeness of our sample. Fourth, because AD and BPH are mostly diseases of aging, we removed anyone who was younger than 40 years old on the index date. Finally, we removed anyone diagnosed with AD on or before the index date.

#### Outcome definition

We defined an individual as having AD if they ever had an ICD-9-CM diagnosis code of 046.11, 046.19, 290.10, 290.11, 290.12, 290.13, 290.20, 290.21, 290.3, 290.40, 290.41, 290.42, 290.43, 291.2, 294.10, 294.11, 331.0, 331.11, 331.19, 331.2, 331.82; or an ICD-10-CM diagnosis code of F01.50, F01.51, F03.90, F03.91, F04, G30.0, G30.1, G30.8, G30.9, G31.1, G31.09, G31.1, G31.81, G31.82, G31.83, G31.84, G31.85, G31.89, G31.9, I69.90, I69.910, I69.911, I69.913, I69.914, I69.915, I69.918, I69.919, I69.920, I69.921, I69.922, I69.923, I69.928, I69.931, I69.932, I69.933, I69.934, I69.939, I69.941, I69.942, I69.943, I69.962, I69.963, I69.964, I69.965, I69.969, I69.990, I69.991, I69.992, I69.993, I69.998, R41.0, R41.1, R41.2, R41.3, R41.4, R41.81, R41.82, R41.83, R41,840, R41.842, R41.843, R41.844, R41.89, R41.9. These codes were based on prior published validations of these code sets to identify AD. For our outcome, we used the first observed date in any setting—outpatient, inpatient, or facilities claim—with one or more of these diagnosis codes.

#### Statistical analysis

We followed individuals from the first observed drug dispensing date (index date) until they were diagnosed with AD or left the Marketscan database, whichever happened first. If an individual left the Marketscan database for any reason, we censored their follow-up on the last observed enrollment date. To reduce confounding, we used inverse probability of treatment weighting (IPTW). IPTW weights each observation according to the inverse probability of the observed treatment outcome—that is, treated individuals have a weight of 1 / the probability of being treated, while untreated individuals have a weight of 1 / the probability of not being treated. After applying IPTW weights, a pseudo-population is derived, for which the values of the confounders used to calculate the treatment probability are more similar. Compared to propensity score matching, a related technique for addressing confounding, IPTW is less likely to create bias and is more data efficient, as it does not discard “unmatched” observations (Austin & Stuart, 2015).

We used entropy balancing to calculate IPTW weights (Hainmueller, 2012). Entropy balancing calculates weights that minimize the negative entropy of the weights, resulting in the exact balance of the means of the included predictor variables. Unlike using a logistic regression model to estimate the weights, entropy balancing is more flexible and provides a closer-to-perfect balance. Additionally, entropy balancing is able to handle more than two classes, an important consideration given we have three groups (TZ/DZ/AZ, tamsulosin, and 5ARI). We used the following classes for entropy balancing: age; drug start year; average number of outpatient encounters per year during the lookback period; whether the patient had any hospitalizations during the lookback period; the number of inpatient encounters per year during the lookback period; the mean number of diagnoses per encounter date during the lookback period; whether the patient had PSA measured during the lookback period; whether the patient had a diagnosis of abnormal PSA; whether the patient had a diagnosis of BPH, slow urinary stream, orthostatic hypotension, or any other hypotension; whether the patient had a uroflow or cystometrogram collected during the lookback period; and the 29 AHRQ-Elixhauser comorbidity flags (van Walraven et al., 2009). The large sample size yields extremely high power to detect clinically negligible differences and the resulting test statistics and p-values are largely uninformative (e.g., p < 10^−16^) regardless of the differences between the groups. We report the standardized mean difference (SMD), also known as Cohen’s d, as our measure of balance. The SMD is the difference between the groups on a particular variable divided by the standard deviation of that measure. Unlike the standard error, the standard deviation does not decrease simply by adding more observations. Common rules of thumb are that SMD with an absolute value of less than 0.1 are balanced, 0.2 are small differences, 0.4 are medium differences, and 0.6 are large differences.

We visualized differences in survival functions using the weighted Kaplan-Meier estimator. To quantify these differences, we estimated hazard ratios (HR) using weighted Cox proportional hazards regression. Because the standard errors of the Cox model do not account for the Kaplan-Meier weightings, we estimated standard errors, test statistics, p-values, and 95% CIs using fractional weighted bootstrapping with 999 bootstrap replicates. We assessed the validity of the proportional hazards using the correlation between the Schoenfeld residuals and time. Additionally, we estimated a model with time-varying coefficients at 1, 3, 5, 7.5, and 10 years. Kaplan-Meier weights were calculated using the R packages *ebal* and *WeightIt*; survival analysis was done using the R package *survival*; all analyses were done using R 4.1.

#### Sensitivity analyses

We conducted several sensitivity analyses:

1. When calculating IPTW weights, extremely large weights can be calculated if the probability of the observed outcome is very small. This can result in a small number of observations having an oversized effect on the outcome. To combat this concern, we repeated our main analysis but trimmed our weights at the 2.5 and 97.5 percentiles, replacing more extreme observed treatment probabilities with the 2.5 and 97.5 percentiles.
2. We investigated whether excluding additional neurological diagnoses affected our results. There may be a period of time before the diagnosis of AD when there is clinical suspicion of AD. During this period, other tentative or more descriptive diagnoses, likely neurological in origin, may be applied to the patient. Specifically, we performed a sensitive analysis in which we excluded anyone with a diagnosis matching one of the Elixhauser neurological comorbidities during the lookback period. Additionally, we estimated an association among those with a neurological history. In both cases, the weights were re-estimated including the same factors as in the main model on the sub-cohorts with and without a neurological history. The HR was again estimated using Cox proportional hazards regression.
3. Prescribing may reflect health status evident to the prescriber but not readily captured in our data. Of specific concern is that a provider may elect to start a patient on tamsulosin or 5ARI, over TZ/DZ/AZ, based on some clinical factor that was unobserved in our data but that may be associated with increased risk of AD. Thus, we repeated our analysis, imposing a 1-year delay between the drug start date and the start of follow-up. This 1-year delay in the start of follow-up reduced potential confounding that may have arisen due to treatment selection factors being related to potential near-term AD risk.
4. Finally, we reviewed the role of additional diagnoses related to AD. We created our case definition to identify the most possible cases of AD. However, many of these AD cases may have been misdiagnosed or assigned as a placeholder diagnosis during a longer neurological course of care. Thus, we performed a sensitivity analysis in which cases of AD were required to include at least two diagnoses of AD, with at least 365 days between the first observed and last observed diagnosis date. We reasoned that when an AD diagnosis recurs, and more specifically, recurs over a long time span, it is more likely that a person had AD, as opposed to another non-AD disorder. If a person was diagnosed with AD but did not meet this heightened case definition, we censored their follow-up on the first observed diagnosis date.

## ACKNOWLEDGMENTS

This work was funded by NIH R01s NS120987 to NSN, and funded by NIH R03 AG078787 and R01 AG086396 to QZ. We gratefully acknowledge Heather Widmayer of the Scientific Editing and Research Communication Core at the University of Iowa Carver College of Medicine and Michael Welsh for critical reading of the manuscript.

## DATA AVAILABILITY

All code and raw data are available at https://narayanan.lab.uiowa.edu.

## AUTHOR CONTRIBUTIONS

QZ, GMA, MAW, and NSN designed the animal experiments. JES and NSN designed the human AD database analysis. QZ, BK, MAW, MO, TL, AB, and RT performed all the animal experiments. DM and ML performed the immunohistochemistry analysis. QZ, TL, MAW, MO and BK maintained and delivered TZ. QZ, JS, JES, and NSN performed all statistical analyses. QZ, JS, JES, GA, and NSN wrote the manuscript, and all authors reviewed and revised the manuscript. JF and SS designed experiments using the yeast system. SS, TH, GP and ML conducted the yeast experiments including microscopy, biochemistry and Western analysis. TH, GP, JF and ML performed the data and statistical analysis, and TH prepared the figures.

## Supplementary Figures

**Figure S1.**
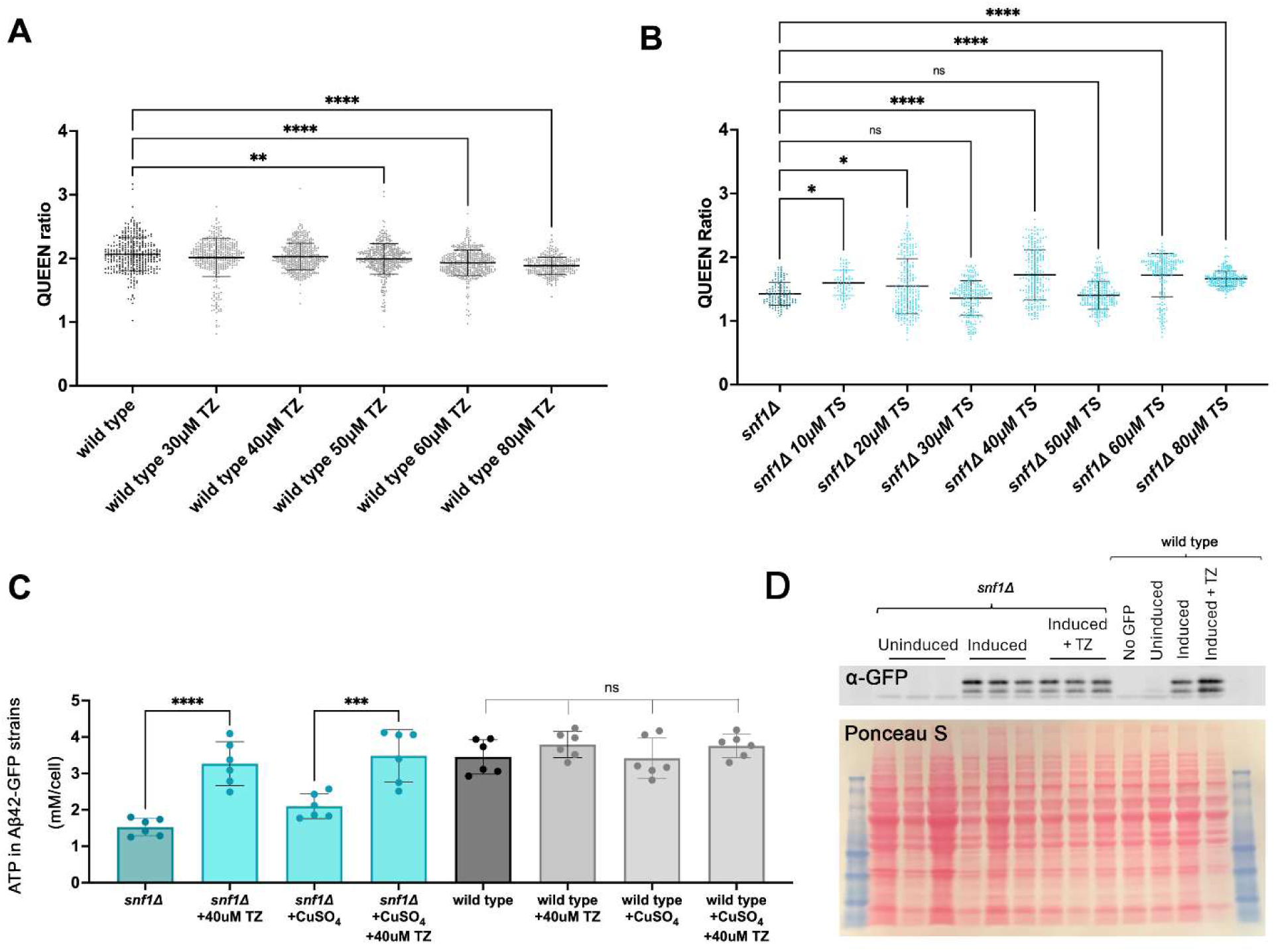
**(A)** Terazosin (TZ) does not raise ATP levels in a wild-type yeast strain. Each data point represents the QUEEN ratio of an individual cell (n=500 cells per dose). Significant changes in the QUEEN ratios at higher levels of TZ reflect a reduction, rather than an increase in ATP and may be indicative of mild toxicity. **(B)** Tamsulosin (TS) does not affect ATP levels in a dose dependent manner. **(C)** Biochemical analysis of ATP showing that the CuSO_4_ used to induce Aβ42-GFP expression had no effect on ATP levels in wild-type or the *snf1* mutant; (n=2). Portions of this analysis are also shown in Fig. 1F. **(D)** Representative α-GFP Western blot analysis showing that the GFP levels are stable regardless of genotype and TZ treatment (n=4; p=0.98). Quantification was performed with reference to the Ponceau S total protein stain shown at the bottom. Significance for panels A-C was determined by ANOVA analysis with a post-hoc Tukey test (****, p<0.0001; ***, p< 0.001; **, p<0.01; *, p<0.05; ns, non-significant).

**Fig S2.**
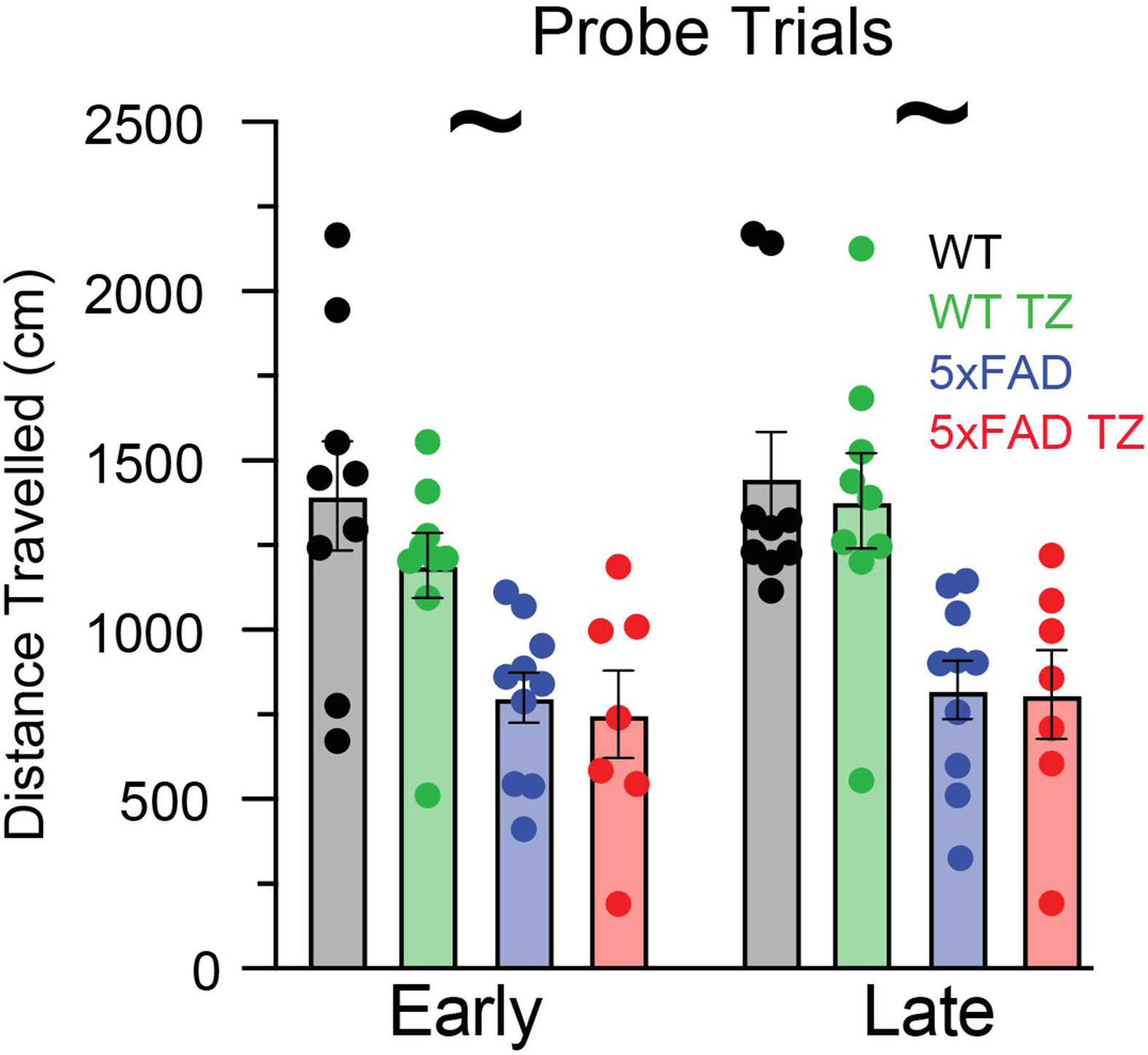
Total distance traveled in the Barnes maze during probe trials during early (Day 2) and late (Day 3) training. ∼ = a main effect of distance with 5xFAD animals traveling less distance, with no effect of TZ or higher interactions for Early (F = 19.2; p = 0.001) and Late (F = 23.5, p = 0.00005) sessions.

**Table S1:** Sample size by exclusion criteria.

|  | **TZ/DZ/AZ** | **Tamsulosin** | **5ARI** |
| --- | --- | --- | --- |
| **Users with 1+ Dispensing Events** | 1,028,561 | 2,854,989 | 653,508 |
| **Exclusive User** | 763,320 | 2,378,238 | 387,203 |
| **New User (At Least 365 Days of Lookback)** | 299,956 | 1,490,963 | 169,360 |
| **One or More Days of Follow-Up** | 299,764 | 1,489,812 | 169,212 |
| **Exclude Female Users** | 225,754 | 1,178,166 | 156,063 |
| **Exclude Age < 40** | 213,302 | 1,046,722 | 123,324 |
| **Exclude AD Case at Baseline** | 209,146 | 1,013,108 | 120,431 |

**Table S2:** Baseline characteristics of new users at first observed claim for study drug.

|  | **Mean (SD)** |  |  | **Standardized Mean Differences - Unadjusted** |  |  |
| --- | --- | --- | --- | --- | --- | --- |
|  | **AZ/DZ/TZ** | **Tamsulosin** | **5ARI** | **AZ/DZ/TZ vs. Tamsulosin** | **AZ/DZ/TZ versus 5ARI** | **Tamsulosin versus 5ARI** |
| **N** | 209,222 | 1,013,108 | 120,537 |  |  |  |
| **Lookback (Years)** | 3.96 (3.06) | 4.87 (3.65) | 4.11 (3.07) |  |  |  |
| **Follow-Up (Years)** | 2.84 (2.80) | 2.73 (2.66) | 2.93 (2.82) |  |  |  |
| **Age** | 61.96 (11.06) | 60.19 (11.16) | 61.46 (12.01) | 0.1560 | 0.0443 | 0.1117 |
| **Year** |  |  |  |  |  |  |
| **2002** | 5,608 (2.7%) | 12,623 (1.2%) | 1,423 (1.2%) | 0.0143 | 0.0150 | 0.0007 |
| **2003** | 5,335 (2.5%) | 15,889 (1.6%) | 2,216 (1.8%) | 0.0098 | 0.0071 | 0.0027 |
| **2004** | 15,915 (7.6%) | 17,377 (1.7%) | 3,019 (2.5%) | 0.0589 | 0.0510 | 0.0079 |
| **2005** | 16,477 (7.9%) | 26,515 (2.6%) | 5,106 (4.2%) | 0.0526 | 0.0364 | 0.0162 |
| **2006** | 11,048 (5.3%) | 30,965 (3.1%) | 5,853 (4.9%) | 0.0222 | 0.0042 | 0.0180 |
| **2007** | 12,980 (6.2%) | 37,256 (3.7%) | 7,479 (6.2%) | 0.0253 | 0.0000 | 0.0253 |
| **2008** | 16,605 (7.9%) | 43,943 (4.3%) | 9,951 (8.3%) | 0.0360 | 0.0032 | 0.0392 |
| **2009** | 20,216 (9.7%) | 58,552 (5.8%) | 13,589 (11.3%) | 0.0388 | 0.0161 | 0.0549 |
| **2010** | 17,659 (8.4%) | 63,388 (6.3%) | 12,054 (10.0%) | 0.0218 | 0.0156 | 0.0374 |
| **2011** | 14,033 (6.7%) | 75,491 (7.5%) | 10,990 (9.1%) | 0.0074 | 0.0241 | 0.0167 |
| **2012** | 14,178 (6.8%) | 87,533 (8.6%) | 9,557 (7.9%) | 0.0186 | 0.0115 | 0.0071 |
| **2013** | 11,884 (5.7%) | 77,878 (7.7%) | 7,484 (6.2%) | 0.0201 | 0.0053 | 0.0148 |
| **2014** | 9,565 (4.6%) | 77,843 (7.7%) | 6,527 (5.4%) | 0.0311 | 0.0084 | 0.0227 |
| **2015** | 7,740 (3.7%) | 69,478 (6.9%) | 5,418 (4.5%) | 0.0316 | 0.0080 | 0.0236 |
| **2016** | 6,306 (3.0%) | 65,490 (6.5%) | 4,413 (3.7%) | 0.0345 | 0.0065 | 0.0280 |
| **2017** | 5,152<br>(2.5%) | 54,700<br>(5.4%) | 3,531<br>(2.9%) | 0.0294 | 0.0047 | 0.0247 |
| **2018** | 4,728<br>(2.3%) | 49,424<br>(4.9%) | 3,187<br>(2.6%) | 0.0262 | 0.0038 | 0.0223 |
| **2019** | 5,078<br>(2.4%) | 53,240<br>(5.3%) | 3,098<br>(2.6%) | 0.0283 | 0.0014 | 0.0268 |
| **2020** | 4,145<br>(2.0%) | 46,044<br>(4.5%) | 2,545<br>(2.1%) | 0.0256 | 0.0013 | 0.0243 |
| **2021** | 4,570<br>(2.2%) | 49,479<br>(4.9%) | 3,097<br>(2.6%) | 0.0270 | 0.0039 | 0.0231 |
| **Outpatient Encounters Per Year** | 20.75<br>(25.70) | 22.50<br>(25.79) | 20.00<br>(22.93) | 0.0702 | 0.0304 | 0.1006 |
| **Ever Hospitalized** | 54,152<br>(25.9%) | 300,543<br>(29.7%) | 25,951<br>(21.5%) | 0.0378 | 0.0435 | 0.0814 |
| **Inpatient Encounters Per Year** | 1.13 (3.52) | 1.23 (3.67) | 0.79<br>(2.90) | 0.0293 | 0.1004 | 0.1296 |
| **Mean Number of Diagnoses Per Encounter Date** | 1.82 (0.84) | 2.15 (1.01) | 1.81<br>(0.77) | 0.3714 | 0.0180 | 0.3894 |
| **Elixhauser Comorbidities** |  |  |  |  |  |  |
| **Cardiac Arrhythmias** | 37,723<br>(18.0%) | 205,749<br>(20.3%) | 21,740<br>(18.0%) | 0.0228 | 0.0001 | 0.0227 |
| **Hypertension** | 144,069<br>(68.9%) | 594,329<br>(58.7%) | 63,780<br>(52.9%) | 0.1020 | 0.1595 | 0.0575 |
| **HTN with Complications** | 25,132<br>(12.0%) | 88,126<br>(8.7%) | 8,896<br>(7.4%) | 0.0331 | 0.0463 | 0.0132 |
| **Neurological Disease** | 8,867<br>(4.2%) | 50,473<br>(5.0%) | 4,808<br>(4.0%) | 0.0074 | 0.0025 | 0.0099 |
| **Diabetes** | 57,921<br>(27.7%) | 248,343<br>(24.6%) | 23,294<br>(19.3%) | 0.0307 | 0.0836 | 0.0529 |
| **Metastatic Cancer** | 2,884<br>(1.4%) | 22,544<br>(2.2%) | 1,683<br>(1.4%) | 0.0085 | 0.0002 | 0.0083 |
| **Solid Tumor** | 22,776<br>(10.9%) | 139,003<br>(13.7%) | 15,073<br>(12.5%) | 0.0283 | 0.0162 | 0.0122 |
| **COPD** | 41,925<br>(20.0%) | 224,732<br>(22.2%) | 21,875<br>(18.1%) | 0.0214 | 0.0189 | 0.0403 |
| **Depression** | 24,019<br>(11.5%) | 144,448<br>(14.3%) | 14,144<br>(11.7%) | 0.0278 | 0.0025 | 0.0252 |
| **Valvular Disease** | 23,263<br>(11.1%) | 120,527<br>(11.9%) | 13,611<br>(11.3%) | 0.0078 | 0.0017 | 0.0060 |
| **Renal Failure** | 21,543<br>(10.3%) | 65,964<br>(6.5%) | 5,954<br>(4.9%) | 0.0379 | 0.0536 | 0.0157 |
| **Psychoses** | 2,339<br>(1.1%) | 14,219<br>(1.4%) | 1,186<br>(1.0%) | 0.0029 | 0.0013 | 0.0042 |
| **Heart Failure** | 18,555<br>(8.9%) | 85,495<br>(8.4%) | 8,927<br>(7.4%) | 0.0043 | 0.0146 | 0.0103 |
| **Peripheral Vascular Disease** | 24,170<br>(11.6%) | 119,164<br>(11.8%) | 12,184<br>(10.1%) | 0.0021 | 0.0144 | 0.0165 |
| **Diabetes with Complications** | 23,763<br>(11.4%) | 101,787<br>(10.0%) | 7,858<br>(6.5%) | 0.0131 | 0.0484 | 0.0353 |
| **Rheumatoid Arthritis** | 9,280<br>(4.4%) | 55,231<br>(5.5%) | 5,044<br>(4.2%) | 0.0102 | 0.0025 | 0.0127 |
| **Obesity** | 23,646<br>(11.3%) | 146,625<br>(14.5%) | 8,742<br>(7.3%) | 0.0317 | 0.0405 | 0.0722 |
| **Fluid and Electrolyte Disorders** | 23,681<br>(11.3%) | 118,996<br>(11.7%) | 9,231<br>(7.7%) | 0.0043 | 0.0366 | 0.0409 |
| **Deficiency Anemia** | 11,700<br>(5.6%) | 58,554<br>(5.8%) | 5,870<br>(4.9%) | 0.0019 | 0.0072 | 0.0091 |
| **Hypothyroidism** | 18,910<br>(9.0%) | 105,013<br>(10.4%) | 11,820<br>(9.8%) | 0.0133 | 0.0077 | 0.0056 |
| **Liver Disease** | 14,741<br>(7.0%) | 104,508<br>(10.3%) | 7,754<br>(6.4%) | 0.0327 | 0.0061 | 0.0388 |
| **Peptic Ulcer Disease** | 3,316<br>(1.6%) | 18,652<br>(1.8%) | 1,632<br>(1.4%) | 0.0026 | 0.0023 | 0.0049 |
| **Blood Loss Anemia** | 2,545<br>(1.2%) | 13,846<br>(1.4%) | 1,335<br>(1.1%) | 0.0015 | 0.0011 | 0.0026 |
| **Weight Loss** | 6,159<br>(2.9%) | 41,391<br>(4.1%) | 3,638<br>(3.0%) | 0.0114 | 0.0007 | 0.0107 |
| **Pulmonary Circulatory Disorders** | 4,355<br>(2.1%) | 26,650<br>(2.6%) | 2,421<br>(2.0%) | 0.0055 | 0.0007 | 0.0062 |
| **Coagulopathy** | 6,396<br>(3.1%) | 39,889<br>(3.9%) | 3,658<br>(3.0%) | 0.0088 | 0.0002 | 0.0090 |
| **Alcohol Abuse** | 5,388<br>(2.6%) | 28,710<br>(2.8%) | 2,174<br>(1.8%) | 0.0026 | 0.0077 | 0.0103 |
| **HIV** | 887 (0.4%) | 4,012<br>(0.4%) | 852<br>(0.7%) | 0.0003 | 0.0028 | 0.0031 |
| **Paralysis** | 2,350<br>(1.1%) | 13,456<br>(1.3%) | 1,058<br>(0.9%) | 0.0020 | 0.0025 | 0.0045 |
| **Lymphoma** | 2,085<br>(1.0%) | 12,771<br>(1.3%) | 1,243<br>(1.0%) | 0.0026 | 0.0003 | 0.0023 |
| **Drug Abuse** | 3,460<br>(1.7%) | 19,260<br>(1.9%) | 1,242<br>(1.0%) | 0.0025 | 0.0062 | 0.0087 |
| **BPH Specific Diagnoses, Procedures** |  |  |  |  |  |  |
| **BPH**<br>**Diagnosis** | 74,826<br>(35.8%) | 367,931<br>(36.3%) | 54,825<br>(45.5%) | 0.0055 | 0.0972 | 0.0917 |
| **Abnormal**<br>**PSA Diagnosis** | 24,023<br>(11.5%) | 134,718<br>(13.3%) | 32,999<br>(27.4%) | 0.0182 | 0.1589 | 0.1408 |
| **PSA**<br>**Measured** | 148,207<br>(70.8%) | 718,142<br>(71.0%) | 87,584<br>(72.7%) | 0.0015 | 0.0182 | 0.0168 |
| **Slow Stream**<br>**Diagnosis** | 7,773<br>(2.8%) | 29,540<br>(2.9%) | 3,105<br>(2.6%) | 0.0016 | 0.0018 | 0.0034 |
| **Uroflow**<br>**Procedure** | 21,715<br>(10.4%) | 105,615<br>(10.4%) | 12,827<br>(10.6%) | 0.0005 | 0.0026 | 0.0022 |
| **Cystometrogram**<br>**Procedure** | 2,875<br>(1.4%) | 12,325<br>(1.2%) | 1,348<br>(1.1%) | 0.0016 | 0.0026 | 0.0010 |
| **Drug Specific**<br>**Factors** |  |  |  |  |  |  |
| **Orthostatic**<br>**Hypotension**<br>**Diagnosis** | 1,678<br>(0.8%) | 9,797<br>(1.0%) | 1,150<br>(1.0%) | 0.0017 | 0.0015 | 0.0001 |
| **Other**<br>**Hypotension**<br>**Diagnosis** | 3,687<br>(1.8%) | 21,784<br>(2.2%) | 2,105<br>(1.7%) | 0.0039 | 0.0002 | 0.0040 |

**Table S3.** Baseline characteristics of weighted pseudo-population.

|  | **AZ/DZ/TZ** | **Tamsulosin** | **5ARI** |
| --- | --- | --- | --- |
| **Effective Sample Size** | 143,949.2 | 982,281.2 | 82,725.3 |
| **Range of Weights**<br>**(narrower is better)** | 0.0832 – 17.9876 | 0.3731 – 2.2825 | 0.0991 – 49.3096 |
| **Coefficient of**<br>**Variability of Weights**<br>**(smaller is better)** | 0.673 | 0.676 | 0.177 |
| **Lookback (Years)** | 4.47 (3.47) | 4.71 (3.54) | 4.41 (3.39) |
| **Follow-Up (Years)** | 2.55 (2.51) | 2.83 (2.77) | 2.68 (2.67) |
| **Age** | 60.57 (10.73) | 60.57 (11.23) | 60.57 (12.18) |
| **Year** |  |  |  |
| **2002** | 3,062.1 (1.5%) | 14,827.7 (1.5%) | 1,764.2 (1.5%) |
| **2003** | 3,652 (1.7%) | 17,684 (1.7%) | 2,104 (1.7%) |
| **2004** | 5,657.3 (2.7%) | 27,394.3 (2.7%) | 3,259.3 (2.7%) |
| **2005** | 7,493.8 (3.6%) | 36,111.9 (3.6%) | 4,296.5 (3.6%) |
| **2006** | 7,457.6 (3.6%) | 36,111.9 (3.6%) | 4,296.5 (3.6%) |
| **2007** | 8,992.1 (4.3%) | 43,542.3 (4.3%) | 5,180.6 (4.3%) |
| **2008** | 10,983.9 (5.2%) | 53,187 (5.2%) | 6,328.1 (5.2%) |
| **2009** | 14,389.4 (6.9%) | 69,677.5 (6.9%) | 8,290.1 (6.9%) |
| **2010** | 14,505.4 (6.9%) | 70,238.8 (6.9%) | 8,356.8 (6.9%) |
| **2011** | 15,660.3 (7.5%) | 75,831.4 (7.5%) | 9,022.2 (7.5%) |
| **2012** | 17,335.8 (8.3%) | 83,944.6 (8.3%) | 9,987.5 (8.3%) |
| **2013** | 15,151.2 (7.2%) | 73,365.9 (7.2%) | 8,728.9 (7.2%) |
| **2014** | 14,635.3 (7.0%) | 70,868 (7.0%) | 8,431.7 (7.0%) |
| **2015** | 12,874.9 (6.2%) | 62,343.6 (6.2%) | 7,417.5 (6.2%) |
| **2016** | 11,873.6 (5.7%) | 57,494.9 (5.7%) | 6,840.6 (5.7%) |
| **2017** | 9,875.2 (4.7%) | 47,818.5 (4.7%) | 6,689 (4.7%) |
| **2018** | 8,933.6 (4.3%) | 43,258.6 (4.3%) | 5,146.8 (4.3%) |
| **2019** | 9,568.8 (4.6%) | 46,334.5 (4.6%) | 5,512.8 (4.6%) |
| **2020** | 8,216.1 (3.9%) | 39,784.5 (3.9%) | 4,733.5 (3.9%) |
| **2021** | 8,903.5 (4.3%) | 43,113.0 (4.3%) | 5,129.5 (4.3%) |
| **Outpatient Encounters**<br>**Per Year** | 22.00 (26.54) | 22.00 (25.41) | 22.00 (24.12) |
| **Ever Hospitalized** | 59,305.6 (28.3%) | 287,173.3 (28.3%) | 34,167.1 (28.3%) |
| **Inpatient Encounters**<br>**Per Year** | 1.18 (3.55) | 1.18 (3.58) | 1.18 (4.31) |
| **Mean Number of**<br>**Diagnoses Per**<br>**Encounter Date** | 2.07 (1.09) | 2.07 (1.19) | 2.07 (0.94) |
| **Elixhauser**<br>**Comorbidities** |  |  |  |
| **Cardiac**<br>**Arrhythmias** | 41,320.7 (6.7%) | 200,085.6 (6.7%) | 23,805.7 (6.7%) |
| **Hypertension** | 124,913.2 (59.7%) | 605,192.5 (59.7%) | 71,909.3 (59.7%) |
| **Neurological Disease** | 9,990.5 (4.8%) | 48,403 (4.8%) | 5,751.3 (4.8%) |
| **Diabetes** | 51,472.5 (24.6%) | 249,379.2 (24.6%) | 29,631.4 (24.6%) |
| **Metastatic Cancer** | 4,221.6 (2.0%) | 20,453.2 (2.0%) | 2,430.3 (2.0%) |
| **Solid Tumor** | 27,538.5 (13.2%) | 133,421.3 (13.2%) | 15,853.2 (13.2%) |
| **COPD** | 44,954.0 (7.3%) | 217,679.1 (7.3%) | 25,898.9 (7.3%) |
| **Depression** | 28,451.2 (4.6%) | 137,768.4 (4.6%) | 16,391.3 (4.6%) |
| **Valvular Disease** | 24,505.7 (11.7%) | 118,727.9 (11.7%) | 14,107.3 (11.7%) |
| **HTN with Complications** | 19,020.3 (9.1%) | 92,151.3 (9.1%) | 10,949.5 (9.1%) |
| **Renal Failure** | 14,553.7 (7.0%) | 70,511.2 (7.0%) | 8,378.2 (7.0%) |
| **Psychoses** | 2,763.1 (1.3%) | 13,387.2 (1.3%) | 1,590.7 (1.3%) |
| **Heart Failure** | 17,590.9 (8.4%) | 85,226.4 (8.4%) | 10,126.6 (8.4%) |
| **Peripheral Vascular Disease** | 24,216.2 (11.6%) | 117,325.2 (11.6%) | 13,940.6 (11.6%) |
| **Diabetes with Complications** | 20,776.3 (9.9%) | 100,659.3 (9.9%) | 11,960.4 (9.9%) |
| **Rheumatoid Arthritis** | 10,831.3 (5.2%) | 52,476.5 (5.52%) | 6,235.3 (5.2%) |
| **Obesity** | 27,876.9 (13.3%) | 135,061.0 (13.3%) | 16,048 (13.3%) |
| **Fluid and Electrolyte Disorders** | 23,653 (11.3%) | 114,598.8 (11.3%) | 13,616.7 (11.3%) |
| **Deficiency Anemia** | 11,860.3 (1.9%) | 57,430.7 (1.9%) | 6,833 (1.9%) |
| **Hypothyroidism** | 21,135.8 (10.1%) | 102,400.9 (10.1%) | 12,167.3 (10.1%) |
| **Liver Disease** | 19,777.4 (9.5%) | 95,819.3 (9.5%) | 11,385.3 (9.5%) |
| **Peptic Ulcer Disease** | 3,674.7 (1.8%) | 17,803.8 (1.8%) | 2,115.5 (1.8%) |
| **Blood Loss Anemia** | 2,761.8 (0.4%) | 13,373.1 (0.4%) | 1,591.1 (0.4%) |
| **Weight Loss** | 7,971.1 (3.8%) | 38,619.2 (3.8%) | 4,588.7 (3.8%) |
| **Pulmonary Circulatory Disorders** | 5,203.7 (2.5%) | 25,211.6 (2.5%) | 2,995.7 (2.5%) |
| **Coagulopathy** | 7,781.2 (1.3%) | 37,678.8 (1.3%) | 25,898.9 (7.3%) |
| **Alcohol Abuse** | 5,651.3 (0.9%) | 27,364.9 (0.9%) | 3,255.8 (0.9%) |
| **HIV** | 895.7 (0.4%) | 4,339.6 (0.4%) | 515.6 (0.4%) |
| **Paralysis** | 2,625.2 (1.3%) | 12,718.6 (1.3%) | 1,511.2 (1.3%) |
| **Lymphoma** | 2,506.5 (1.2%) | 12,143.6 (1.2%) | 1,442.9 (1.2%) |
| **Drug Abuse** | 3,731.6 (1.8%) | 18,079.2 (1.8%) | 2,148.2 (1.8%) |
| **BPH Specific Diagnoses, Procedures** |  |  |  |
| **BPH Diagnosis** | 77,477.9 (37.1%) | 375,373.1 (37.1%) | 44,602 (37.1%) |
| **Abnormal PSA Diagnosis** | 29,853.1 (14.3%) | 144,635 (14.3%) | 17,185.6 (14.3%) |
| **PSA Measured** | 148,698.3 (71.1%) | 720,429.0 (71.1%) | 85,601.7 (71.1%) |
| **Slow Stream Diagnosis** | 5,941.7 (2.9%) | 28,980.8 (2.9%) | 3,443.5 (2.9%) |
| **Uroflow Procedure** | 21,824.7 (10.4%) | 105,738.4 (10.4%) | 12,563.9 (10.4%) |
| **Cystometrogram Procedure** | 2,575.9 (1.2%) | 12,480.2 (1.2%) | 1,482.9 (1.2%) |
| **Drug Specific Factors** |  |  |  |
| **Orthostatic Hypotension Diagnosis** | 1,966.3 (0.9%) | 9,526.7 (0.9%) | 1,132 (0.9%) |
| **Other Hypotension Diagnosis** | 4,294 (2.1%) | 20,804 (2.1%) | 2,471.9 (2.1%) |

**Table S4:**
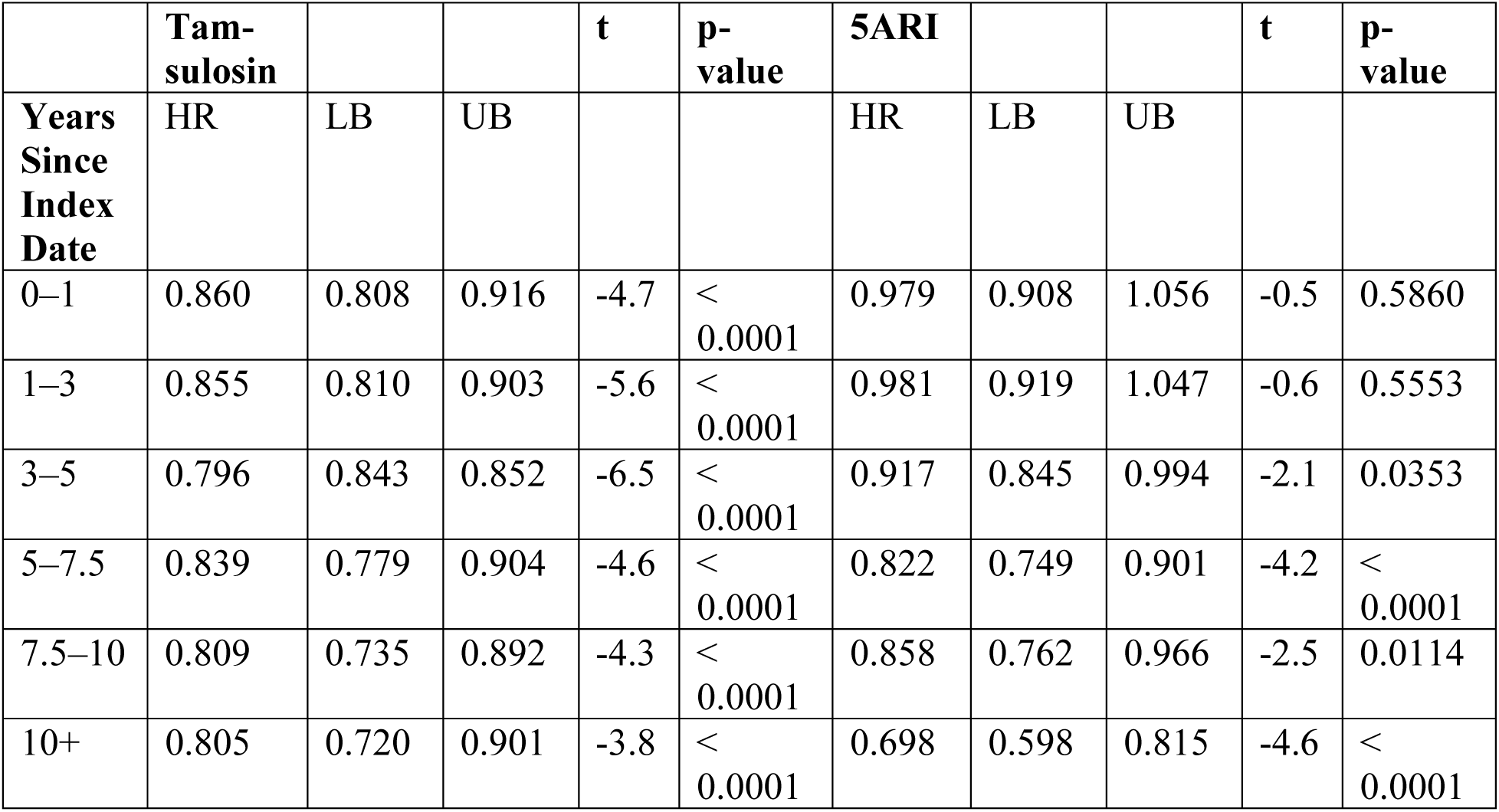
Results of model with time-varying coefficients for TZ/DZ/AZ and 5ARI vs. tamsulosin.

**Table S5:** Results of sensitivity analyses with re-defined cohorts, follow-up delays, and alternative case definitions.

|  | **Trimmed Weights** | **Positive Neurological History** | **No Neurological History** | **One Year Delay in Follow-Up** | **2+ Diagnoses, 365 days** |
| --- | --- | --- | --- | --- | --- |
| **Person-Years at Risk** |  |  |  |  |  |
| **TZ/DZ/AZ** | 512,670 | 19,411 | 514,504 | 521,884 | 533,753 |
| **Tamsulosin** | 2,870,250 | 118,478 | 2,751,358 | 2,683,610 | 2,870,393 |
| **5ARI** | 313,238 | 10,898 | 312,095 | 315,493 | 322,522 |
| **Cases of AD** |  |  |  |  |  |
| **TZ/DZ/AZ** | 6,665 | 1,208 | 5,875 | 5,197 | 1,760 |
| **Tamsulosin** | 47,225 | 8,204 | 39,266 | 34,262 | 13,452 |
| **5ARI** | 4,519 | 689 | 4,058 | 3,576 | 1,311 |
| **Incidence Rate** |  |  |  |  |  |
| **TZ/DZ/AZ** | 13.0 per 1,000 | 62.2 per 1,000 | 11.4 per 1,000 | 10.0 per 1,000 | 3.30 |
| **Tamsulosin** | 16.5 per 1,000 | 69.2 per 1,000 | 14.3 per 1,000 | 12.8 per 1,000 | 4.69 |
| **5ARI** | 14.4 per 1,000 | 63.2 per 1,000 | 13.0 per 1,000 | 11.3 per 1,000 | 4.06 |
| **HR (95% CI) [t, p-value]** |  |  |  |  |  |
| **TZ/DZ/AZ** | 0.806<br>(0.806, 0.806)<br>[-1,091.6, p < 10<sup>-16</sup>] | 0.897<br>(0.895, 0.900)<br>[-87.0, p < 10<sup>-16</sup>] | 0.825<br>(0.825, 0.826)<br>[-952.7, p < 10<sup>-16</sup>] | 0.823<br>(0.823, 0.824)<br>[-969.3, p < 10<sup>-16</sup>] | 0.726<br>(0.726, 0.727)<br>[-1,120.1, p < 10<sup>-16</sup>] |
| **Tamsulosin** | Reference |  |  |  |  |
| **5ARI** | 0.880<br>(0.880, 0.881)<br>[-653.3, p < 10<sup>-16</sup>] | 0.911<br>(0.909, 0.914)<br>[-64.7, p < 10<sup>-16</sup>] | 0.920<br>(0.920, 0.921)<br>[-445.2, p < 10<sup>-16</sup>] | 0.908<br>(0.907, 0.908)<br>[-492.1, p < 10<sup>-16</sup>] | 0.878<br>(0.877, 0.879)<br>[-293.4, p < 10<sup>-16</sup>] |

